# Porcine epidemic diarrhea virus infection promotes Peyer’s patch immune induction and epithelial defense via single-cell transcriptional reprogramming

**DOI:** 10.1101/2025.06.13.659514

**Authors:** Jayne E. Wiarda, Bailey Arruda, Eraldo L. Zanella, Hanjun Kim, Samantha J. Hau, Jianqiang Zhang, Alexandra C. Buckley

**Affiliations:** Virus and Prion Research Unit, National Animal Disease Center, Agricultural Research Service, United States Department of Agriculture, Ames, IA, USA; Oak Ridge Institute for Science and Education, Agricultural Research Service Participation Program, Oak Ridge, TN, USA; Department of Veterinary Diagnostics and Production Animal Medicine, Iowa State University College of Veterinary Medicine, Ames, IA, USA

**Author notes:** Address correspondence to Jayne E. Wiarda,.

## Abstract

Porcine epidemic diarrhea virus (PEDV) is an enteric coronavirus causing gastrointestinal disease in swine. To mitigate PEDV risks to swine health, agricultural economic losses, and global food security, a better understanding of host-pathogen interactions is required. Using single-cell RNA sequencing (scRNA-seq), we studied transcriptional cellular responses to PEDV infection in two anatomical compartments of intestinal jejunum: first-line barrier defenses of epithelia and underlying immune induction in Peyer’s patches. PEDV infection altered gene expression across all cell types and was associated with antiviral response pathways, indicating coordinated transcriptional reprogramming of diverse cell types. Signaling network inferences showed macrophages, dendritic cells, and non-resting B cells had increased signaling to T follicular helper cells associated with processes of T cell-dependent B cell activation during PEDV infection, indicating transcriptional promotion of immune induction in Peyer’s patches. Mature enterocytes were the primary targets for PEDV infection, and a 190-gene signature indicative of antiviral immune defense represented a conserved enterocyte response to PEDV infection, regardless of enterocyte stress state or infection status. The 190-gene signature was specific to the epithelial lineage and escalated with enterocyte maturation, indicating antiviral transcriptional reprogramming is mobilized across matured enterocytes of PEDV-infected intestinal segments. Results exemplify key roles of Peyer’s patches and epithelia as integral components for coordinated antiviral responsiveness in the intestine. Improved understandings of host-pathogen interactions during PEDV infection can identify indicators of PEDV protection versus susceptibility to target for future intervention strategies.

**IMPORTANCE:** Porcine epidemic diarrhea virus (PEDV) causes high death rates in young pigs and production losses in older animals, yet prevention and treatment options for PEDV remain limited. Understanding how PEDV infects pigs and how cells can control infection is crucial for developing better prevention and treatment strategies. Our work explores how intestinal cells are impacted by PEDV infection. We find all intestinal cells responded to infection despite diverse origins and functions. Processes promoting immune responses and epithelial barrier defenses against PEDV were initiated, culminating in a highly coordinated and conserved antiviral response. Results identify targets that may be useful in developing new PEDV prevention and control strategies that could positively impact animal health, agricultural economic prosperity, and global food security.

## INTRODUCTION

Porcine epidemic diarrhea virus (PEDV), the causative agent of porcine epidemic diarrhea (PED), is an enteric coronavirus causing gastrointestinal disease in pigs [1]. PEDV has been an economic and animal health burden to the swine industry since its emergence in Europe during the 1970s, and PEDV was introduced to the United States in 2013 [1]. By 2014, widespread losses attributed to PEDV outbreaks cost the United States swine industry an estimated 1.8 billion dollars [2], making PED one of the most economically important swine diseases. PEDV infection causes the highest mortalities in neonates (litter mortality of up to 95% in farrowing operations) [3] but also causes reduced performance at later stages of pig production [4–6]. PEDV continues to be detected in diagnostic cases, including in samples from farrowing and wean-to-finish operations [7], indicating PEDV persists in United States swine of all production stages. However, prevention and treatment options for PEDV still remain limited [1, 8–10].

Deciphering how localized enteric antiviral responses are induced by PEDV could prove crucial for identifying targets for preventive or therapeutic strategies. The epithelial barrier is a first-line defense against enteric pathogens [11, 12] and is the primary target of PEDV infection [13]. Several studies have indicated strategies fortifying epithelial barrier integrity may decrease susceptibility to PEDV [14–17], demonstrating the importance of a healthy epithelium in preventing PEDV infection. Upon infection, PEDV causes breakdown of the epithelial barrier [18–21], leading to reduced nutrient absorption that can potentiate production inefficiencies and promiscuous trans-epithelial passage of other luminal pathogens or molecules that can have harmful effects. When intestinal barrier integrity is compromised, the underlying immune network must act as a second layer of protection. The intestinal immune system plays a key role in combatting PEDV, as the importance of stimulating localized enteric immunity has been demonstrated by immunomodulatory treatments. Orally-administered PEDV vaccines and oral ingestion of PEDV-infected biomaterials provide superior passive lactogenic immunity and protection against later PEDV infection due to induction of localized intestinal immunity [9, 10, 22]. Thus, first-line epithelial defenses and underlying immune networks within the intestinal tract are both key components to protect against PEDV. Despite a basic understanding of local enteric immunity to PEDV, the immune response to PEDV infection has gone severely understudied in Peyer’s patches. Peyer’s patches are the only organized lymphoid tissue embedded directly in the small intestine and do not rely on lymphatic migration of luminally-derived immunogens [23]. Peyer’s patches play critical roles in inducing immune activation versus tolerance to immune stimuli at local intestinal sites, yet the role of Peyer’s patches, and the complex cell networks that comprise them, remain understudied in the context of PEDV immunity.

Herein, we utilize single-cell RNA sequencing (scRNA-seq) to study how the immune landscape of pig jejunum containing Peyer’s patches is impacted by PEDV infection. We focus primarily on two components of intestinal organization: first-line barrier defenses of the intestinal epithelium and underlying immune induction networks in Peyer’s patches. Results indicate diverse intestinal cell types partake in a culminating antiviral transcriptional response that can be further utilized for identifying targets of preventative and therapeutic strategies against PEDV. Thus, understanding dynamics of PEDV infection is critical for promoting swine health and global security of pork as a food source. Though PEDV does not infect humans, inter-species spillovers of coronavirus infections remain a global concern [24], and disease dynamics of PEDV may have cross-implications for study of coronavirus infections in other species, including humans.

## RESULTS

### Experimental PEDV inoculation causes clinical disease and atrophic enteritis in weaned pigs

A basic experimental overview is available in **Figure 1A**. Four-week-old weaned pigs (metadata in **Supplementary File 1**) were orally administered PEDV (n=5) or mock inoculum (n=3). Animals were monitored daily for clinical signs of disease (diarrhea score), and daily rectal swabs were collected and tested for PEDV RNA copies via reverse transcription quantitative polymerase chain reaction (RT-qPCR). Necropsies were performed at 5 days post-inoculation (dpi) to collect jejunum containing Peyer’s patches from each animal. The proximal-most region of jejunum containing Peyer’s patches (proximal positioning of tissues shown in **Supplementary Figure 1A**) was partitioned into separate pieces allocated for (1) cell isolations used for scRNA-seq, (2) RNAlater storage for PEDV transcript detection via RT-qPCR, and (3) formalin fixative for histologic evaluation. Tissue for cell isolations and RNAlater was collected to include only areas containing Peyer’s patches overlaid with intestinal mucosa, while tissue collected for histopathology contained an entire transverse section of jejunum including areas with and without Peyer’s patches (**Supplementary Figure 1B**).

**Figure 1.**
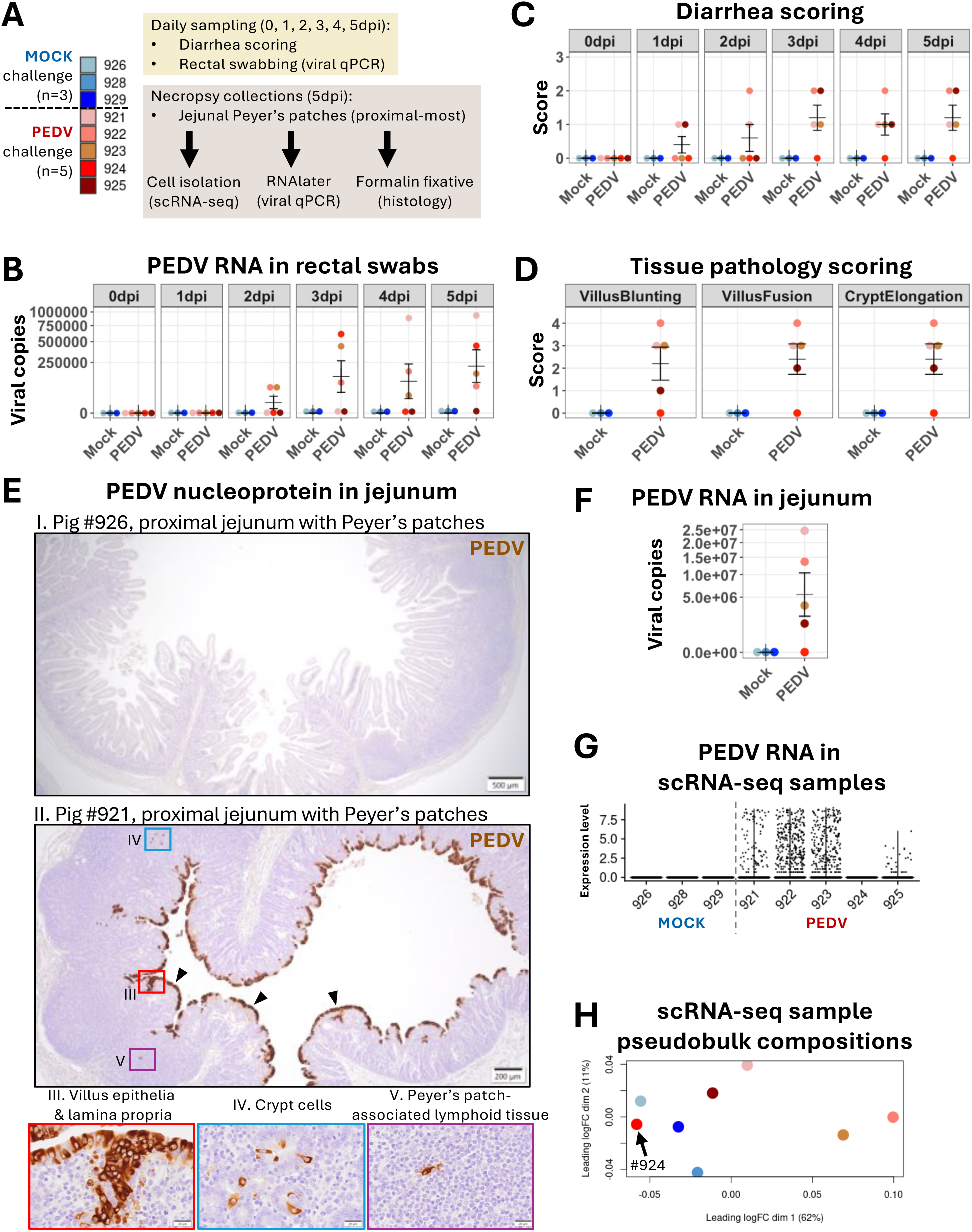
Experimental PEDV inoculation established successful infection in weaned pigs. **(A)** Experimental overview showing pigs inoculated with PEDV (n=5) or mock (n=3) inoculum and samples taken daily (top box) or at 5dpi necropsy (bottom box). The color key for individual animal identities is applied to panels throughout the figure (panels B, C, D, F, H). **(B)** PEDV viral copies (y-axes) from rectal swabs of mock (blue) or PEDV-inoculated (red) pigs (x-axes). **(C)** Diarrhea scores (y-axes) of mock (blue) or PEDV-inoculated (red) pigs (x-axes). **(D)** Atrophic enteritis scores for jejunum sections collected from mock (blue) or PEDV-inoculated (red) pigs at 5dpi, including epithelial villus blunting (left), epithelial villus fusion (center), or epithelial crypt elongation (right). **(E)** Photomicrographs of jejunal tissues stained for PEDV nucleoprotein (brown). I) No PEDV immunolabeling and histologically normal jejunum with Peyer’s patches from mock pig #926; 20X magnification. II) Abundant PEDV immunolabeling and atrophic enteritis (villus blunting, villus fusion, crypt elongation) in jejunum with Peyer’s patches from PEDV-inoculated pig #921; 20X magnification. PEDV nucleoprotein was detected in mature enterocytes (arrowheads and red box), lamina propria (red box), crypt cells (blue box), and Peyer’s patch-associated lymphoid tissue (purple box). III-V) PEDV immunolabeling of boxed regions shown in panel II; 600X magnification. PEDV nucleoprotein was detected in villus enterocytes and cells of lamina propria (III), crypts (IV), and Peyer’s patch-associated lymphoid tissue (V). **(F)** PEDV viral copies (y-axis) obtained from jejunum of mock (blue) or PEDV-inoculated (red) pigs collected at 5dpi. **(G)** PEDV RNA expression (y-axis) detected in individual scRNA-seq animal samples (x-axis). Each dot represents one cell from a respective sample. **(H)** Multidimensional scaling plot of individual scRNA-seq samples based on total library pseudobulk expression profiles. Expression of all detected genes was used to create dimensional reductions. Bar intervals in **B, C, D, F** indicate mean and standard error of the mean. Abbreviations: dim (dimension); dpi (days post-inoculation); H&E (hemotoxylin and eosin); logFC (log fold-change); PEDV (porcine epidemic diarrhea virus); qPCR (quantitative polymerase chain reaction); scRNA-seq (single-cell RNA sequencing)

PEDV infection was confirmed by temporally-increasing PEDV RNA copies detected in rectal swabs of PEDV-inoculated pigs (**Figure 1B**). Diarrhea (**Figure 1C**) and microscopic lesions (**Figure 1D**) were detected in 4 of 5 PEDV-inoculated pigs. Microscopic lesions included villus blunting, villus fusion, and crypt elongation (**Figure 1D** and **1E panels I, II**). Immunohistochemical detection of PEDV nucleoprotein (detected in 4 of 5 PEDV-inoculated pigs) revealed virus localized primarily to villus epithelium (**Figure 1E panel II**), though PEDV staining was also identified in epithelial crypts and non-epithelial locations, including cells within the lamina propria and organized lymphoid tissue associated with Peyer’s patches (**Figure 1E panels III, IV, V**). Presence of PEDV in collected jejunal tissue was further confirmed via quantification of PEDV RNA copies (**Figure 1F**).

PEDV-inoculated pig #924 did not develop diarrhea (**Figure 1C**), did not exhibit microscopic lesions (**Figure 1D**), had minimal PEDV transcript copies detected in jejunal tissue (**Figure 1F**), and did not have immunohistochemical detection of PEDV in jejunal tissue (similar to **Figure 1E panel I**). scRNA-seq revealed no cells from pig #924 contained PEDV RNA, unlike other PEDV-inoculated animals (**Figure 1G**). Comparison of pseudobulk compositions from scRNA-seq samples (**Figure 1H**) revealed the overall transcriptional composition of pig #924 was more similar to mock than other PEDV-inoculated pigs. Although rectal swabs for #924 tested PEDV-positive (**Figure 1B**), indicating PEDV was detected in the intestinal tract, lack of evidence for localized PEDV infection in the jejunal tissue harvested for further analyses and overall transcriptional similarities of pig #924 jejunum to mock animals led to omission of cells from pig #924 from any further comparisons of mock versus PEDV treatment groups.

### PEDV infection skews diverse cell type transcriptomes towards antiviral responsiveness

scRNA-seq of isolated cells from jejunum containing Peyer’s patches of eight pigs resulted in a dataset of 40,522 total cells (**Figure 2A**). Transcriptional profiles of cells were used to create 43 clusters (**Figure 2B**) that were annotated into specific cell types (**Figure 2C**) and cell lineages (**Figure 2D**) based on expression of canonical gene markers (**Supplementary Figure 2A**). Cluster 32 was identified as a low-quality B cell cluster having low gene feature complexity (**Supplementary Figure 2B**) and was consequently removed from further analyses. Remaining cells belonging to all clusters, cell types, and cell lineages were represented in both PEDV and mock samples used for further comparisons (**Supplementary Figure 2C**).

**Figure 2.**
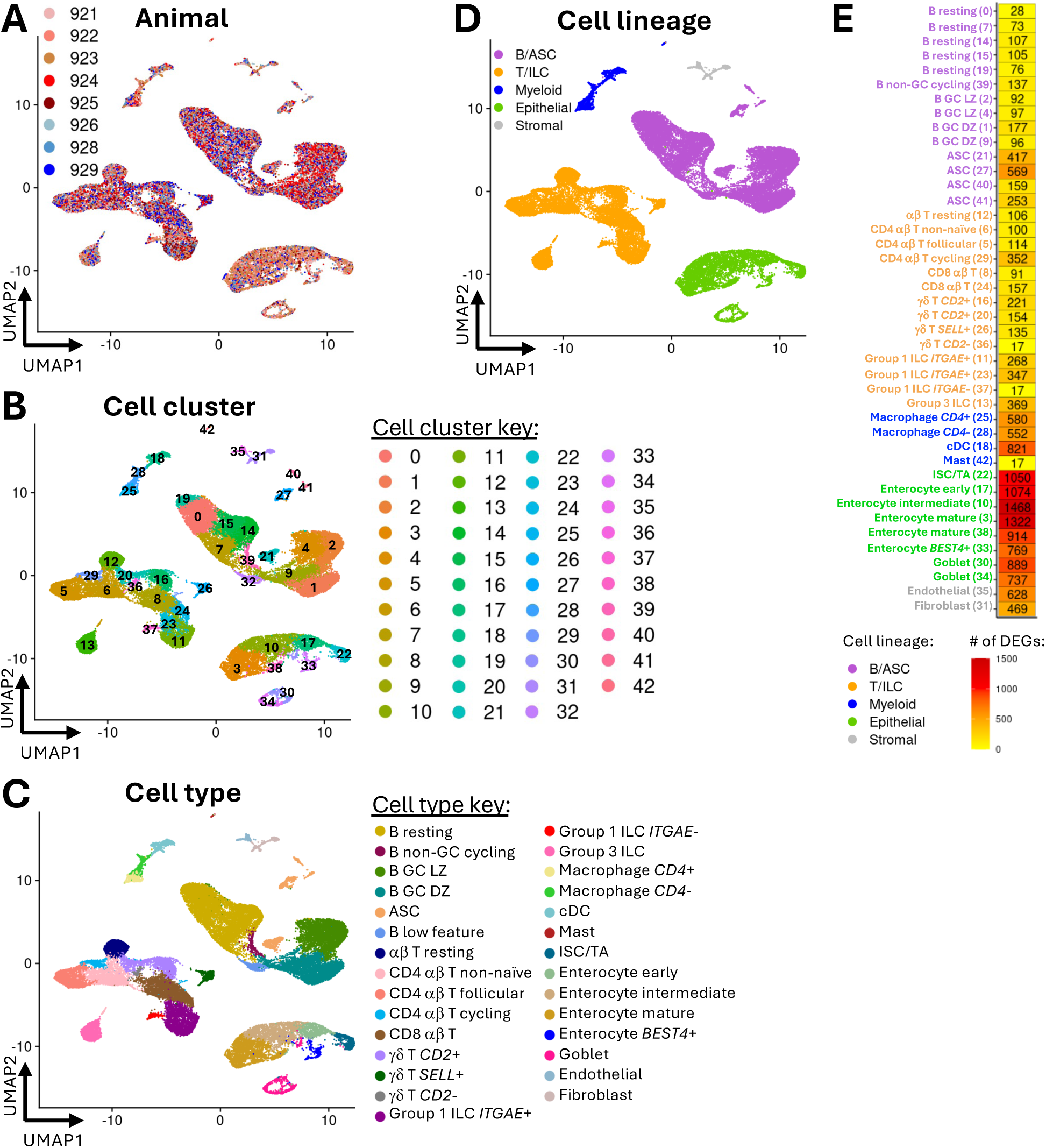
scRNA-seq captures the diverse cell landscape of jejunum containing Peyer’s patches during PEDV or mock infection. **(A-D)** UMAP plots of animal IDs **(A)**, cell clusters **(B)**, cell types **(C)**, and cell lineages **(D)** identified from scRNA-seq data of pig jejunum containing Peyer’s patches. Each dot represents one cell. Dot color corresponds to assignment of a cell to an animal **(A)**, cluster **(B)**, cell type, or cell lineage **(D)**. **(E)** Heatmap showing the number of DEGs (fill color) between mock and PEDV cells within each cell cluster (y-axis). Identified DEGs had corrected p-values <0.05, were expressed in at least 10% of one cell population being compared, and had a |logFC| >0.25. Abbreviations: ASC (antibody-secreting cell); cDC (conventional dendritic cell); DEG (differentially expressed gene); DZ (dark zone); GC (germinal center); ILC (innate lymphoid cell); ISC (intestinal stem cell); logFC (log fold change); LZ (light zone); PEDV (porcine epidemic diarrhea virus); scRNA-seq (single-cell RNA sequencing); TA (transit amplifying); UMAP (uniform manifold approximation and projection)

To understand how PEDV infection impacts specific cell subsets, differential gene expression (DGE) analysis was performed to identify differentially expressed genes (DEGs) between cells derived from mock versus PEDV treatments within each cluster (**Supplementary File 2**). Every cluster contained DEGs (**Figure 2E**), indicating PEDV infection caused transcriptional disruptions across all types of cells. In line with villus epithelium being a primary target of PEDV infection in **Figure 1E**, epithelial cells had the greatest number of DEGs; however, several immune cell subsets also had relatively high numbers of DEGs, including clusters of B cells, antibody-secreting cells (ASCs), CD4 αβ T cells, CD8 αβ T cells, *CD2*+ γδ T cells, group 1 innate lymphoid cells (ILCs), group 3 ILCs, macrophages, and conventional dendritic cells (cDCs) all containing >100 DEGs (**Figure 2E**). The top biological processes enriched in the PEDV group (**Supplementary Figure 3A-E**; **Supplementary File 3**) cumulatively suggested genes supporting antiviral immune processes (e.g. antigen processing, MHC I presentation, type I interferon responsiveness, innate immunity, cytokine production/signaling, viral responsiveness) had increased expression in PEDV compared to mock pigs across various cell types.

### T follicular helper cells are central hubs of upregulated immune network signaling promoting Peyer’s patch immune induction during PEDV infection

Further coordination of cellular responses to PEDV infection were assessed by analyzing signaling networks in mock versus PEDV samples, where increased/decreased signaling strengths between cell types were inferred from higher/lower expression of genes encoding receptors or ligands for cell signaling pathways. For all cell clusters, cumulative incoming and outgoing signals were stronger in the PEDV group (positive axes values shown in **Figure 3A**), indicating transcriptional promotion of cellular communication in response to PEDV infection. Cell clusters with the largest overall increases in incoming/outgoing signaling strengths were generally epithelial, stromal, and myeloid lineage cells, including classical antigen-presenting cell (APC) subsets of macrophages (clusters 25, 28) and cDCs (cluster 18) (**Figure 3A**).

**Figure 3.**
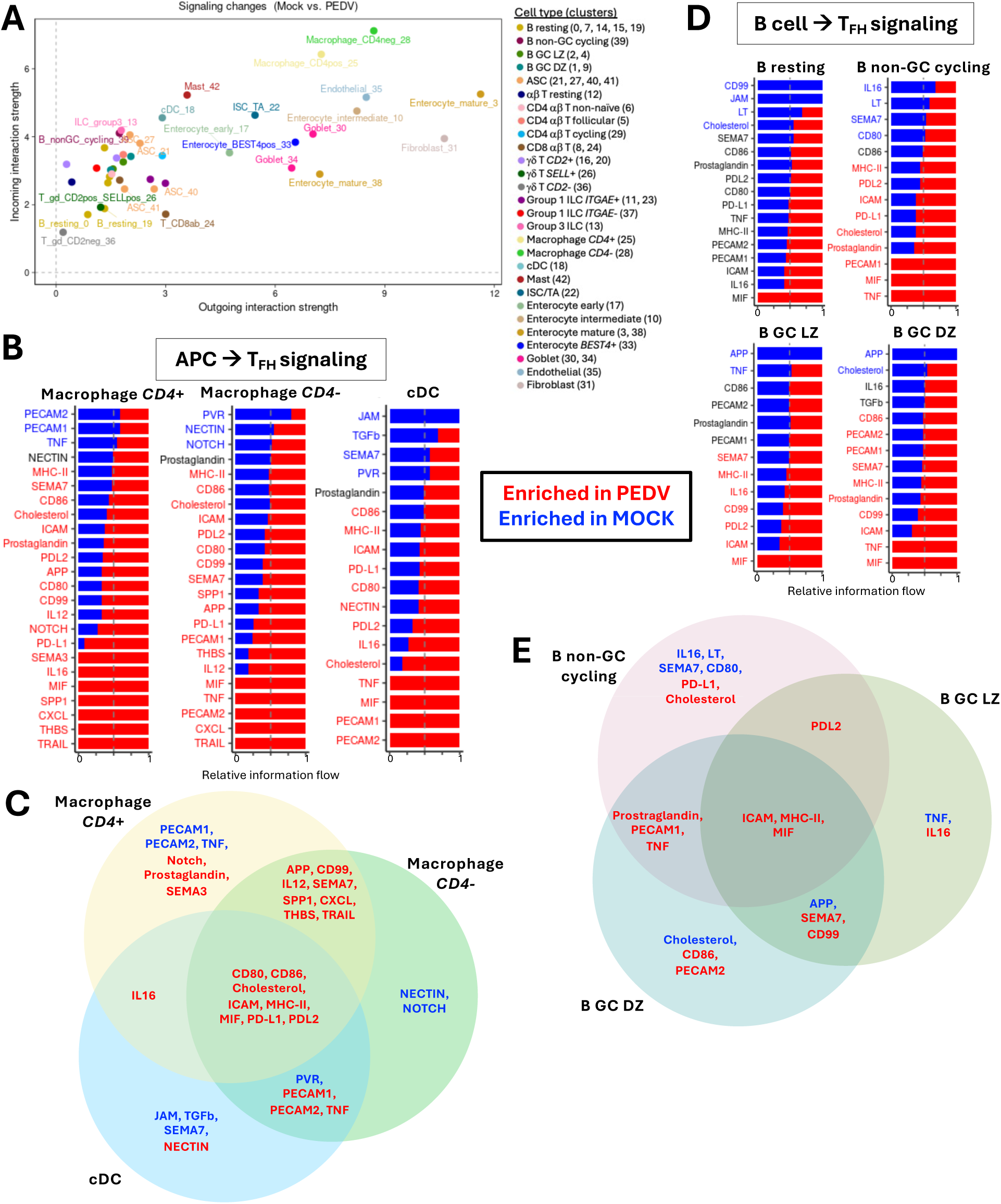
PEDV infection upregulates genes involved in key cell signaling networks. **(A)** Scatter plot showing cumulative incoming (y-axis) and outgoing (x-axis) cell signaling interaction strengths for cell clusters. Positive axis values indicate greater signaling strengths in PEDV relative to mock cells. **(B, D)** Stacked bar plots showing signaling networks (y-axes) with largest relative differences in information flow (x-axes) between mock and PEDV cells. Signaling network names shown in blue were enriched in mock cells, while signaling networks shown in red were enriched in PEDV cells. Plots show signaling networks where signals were received by Tfh cells from various signaling senders (APCs in **(B)** and B cells in **(D)**). Cell types sending signals to Tfh cells are listed above respective plots. **(C, E)** Venn diagrams showing overlap of signaling networks enriched in mock (blue) or PEDV (red) cells. The diagram in **(C)** shows signaling networks sent from APCs to Tfh cells from **(B).** The diagram in **(E)** shows signaling networks sent from non-resting B cells to Tfh cells from **(D).** Abbreviations: APC (antigen presenting cell); ASC (antibody-secreting cell); cDC (conventional dendritic cell); DZ (dark zone); GC (germinal center); ILC (innate lymphoid cell); ISC (intestinal stem cell); LZ (light zone); PEDV (porcine epidemic diarrhea virus); TA (transit amplifying); Tfh (T follicular helper)

For both PEDV and mock groups, macrophages and cDCs were strong signal senders to resting αβ T cells and other CD4 αβ T cell subsets, with the highest overall signaling strengths being detected for signals received by CD4 αβ T follicular helper (Tfh) cells in cluster 5 (**Supplementary Figure 4A**), suggesting APCs act as a signaling source for Tfh cells. Higher relative signaling strengths from macrophage/cDC clusters to Tfh cells were also detected in PEDV over mock samples (**Supplementary Figure 4B**), indicating APC-to-Tfh signaling was strengthened during PEDV infection. Tfh cells require specific APC-derived signaling cues for activation [25, 26]. To determine if PEDV infection influenced APC-derived signaling cues to Tfh cells, we identified specific APC-to-Tfh cell signaling networks that were enriched due to large differences in relative information flows between PEDV and mock samples (**Figure 3B**). For all three macrophage/cDC cell types sending signals to Tfh cells, several signaling networks were enriched in the PEDV group (**Figure 3C**), including networks with known roles in APC activation of T cells (CD80, CD86, ICAM, MHC-II, PD-L1, PD-L2) [25–28].

After Tfh cells have been activated, they can interact with antigen-presenting B cells to facilitate T cell-dependent B cell activation [25, 26, 29]. We next investigated signaling networks between Tfh and B cells, which also involved high-strength interactions for B-to-Tfh cell signaling in both mock and PEDV groups (**Supplementary Figure 4A**). Signaling strengths were higher in the PEDV relative to mock group (**Supplementary Figure 4B**), indicating PEDV infection resulted in strengthened B-to-Tfh signaling. There were no signaling networks from resting B cells to Tfh cells that were enriched in the PEDV group; however, multiple signaling networks from remaining non-resting B cell types to Tfh cells were enriched in the PEDV group (**Figure 3D**). In all three non-resting B cell types sending signals to Tfh cells, ICAM and MHC-II signaling networks were enriched in the PEDV group (**Figure 3E**), which are required for T cell-dependent B cell activation processes [25, 26, 29, 30].

### PEDV preferentially infects mature enterocytes with polarized homeostatic or stressed transcriptional states

Cells directly infected with PEDV were identified to better understand dynamics of host cellular tropism occurring during PEDV infection. PEDV RNA was only detected in cells from the PEDV treatment group (**Figure 4A; Supplementary Figure 5A**) and was used as a readout to infer actively infected cells. Various immune, epithelial, and stromal cell clusters contained PEDV RNA, though the highest percentages of infected cells within PEDV samples were detected in mature enterocyte clusters (26% of PEDV group cells in cluster 3; 50% of PEDV group cells in cluster 38) (**Figure 4A**). Frequent detection of PEDV RNA in mature enterocyte clusters coordinated with immunohistochemical detection of PEDV nucleoprotein primarily localized to villus epithelial cells (**Figure 1E panel II**), while lower percentage detection of PEDV RNA in additional cell types corresponded to less frequent immunohistochemical detection of PEDV in crypts, lamina propria, and Peyer’s patches (**Figure 1E panels II, IV, V)**.

**Figure 4.**
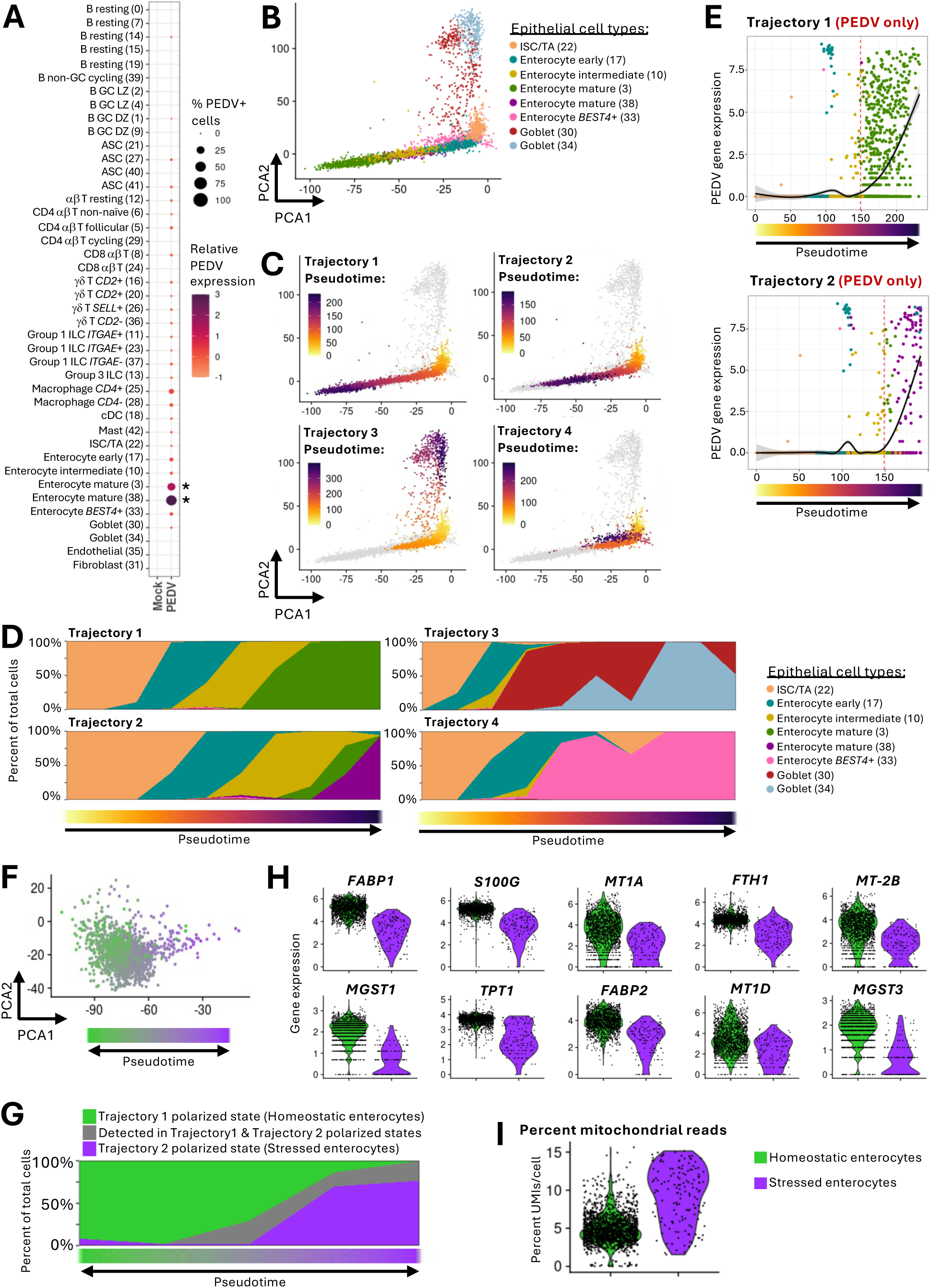
PEDV preferentially infects enterocytes in polarized stress states. **(A)** Dot plot of PEDV RNA expression in mock versus PEDV cells (x-axis) for each cell cluster (y-axis). Dot size indicates the percentage of cells with PEDV RNA; dot color indicates relative average expression level of PEDV RNA, for those cells expressing PEDV RNA within each cluster. Asterix to the right of the plot indicate PEDV RNA was significantly differentially expressed between mock and PEDV groups for indicated clusters. DEGs had corrected p-values <0.05, were expressed in at least 10% of one cell population being compared, and had a |logFC| >0.25. **(B-C)** PCA plots of only epithelial cells. Each dot represents one cell. The color of the dot corresponds to assignment of a cell to a cluster **(B)** or a pseudotime value **(C)** calculated from projected trajectories occurring in the epithelial cell dataset. **(C)** Stacked bar plots showing the percent of cells (y-axes) belonging to each cell cluster (fill colors) across pseudotime values (x-axes) of trajectories shown in **(C)**. **(D)** Scatter plots showing the expression level for PEDV RNA (y-axes) across pseudotime (x-axes) for trajectories 1 (top) and 2 (bottom) shown in **(C-D).** Only cells from PEDV pigs are shown on the plot. Each dot represents one cell. Dot color corresponds to assignment of a cell to a cluster. A red dotted line at x=148.796 indicates a bifurcation of trajectory 1 and 2 and corresponds to a knot value obtained when fitting a generalized additive model to each trajectory. A line of best fit (solid, black) shows PEDV expression levels across pseudotime. **(E)** PCA plot of only cells occurring after the x-intercept knot value (x=148.796) shown in trajectory 1 or 2, as identified in **(E)**. Each dot represents one cell. The color of the dot corresponds to a pseudotime value calculated from a projected trajectory occurring in the data subset. **(F)** Stacked bar plot showing the percent of cells (y-axis) belonging to trajectory 1 or 2 (shown in **(C-D)**; fill color) across pseudotime values (x-axis) of the trajectory shown in **(F)**. Cells identified only in trajectory 1 were called homeostatic enterocytes (green); cells identified only in trajectory 2 were called stressed enterocytes (purple). **(G)** Violin plots of gene expression (y-axes) detected in homeostatic versus stressed enterocytes (x-axes) identified in **(G)**. Each dot represents one cell from a respective sample. The 10 DEGs with highest logFC expression increased in homeostatic versus stressed enterocytes are shown. DEGs had corrected p-values <0.05, were expressed in at least 10% of one cell population being compared, and had a |logFC| >0.25. **(H)** Violin plot of mitochondrial read content (y-axis) detected in homeostatic versus stressed enterocytes (x-axis) identified in **(G)**. Each dot represents one cell from a respective sample. Abbreviations: ASC (antibody-secreting cell); cDC (conventional dendritic cell); DEG (differentially expressed gene); DZ (dark zone); GC (germinal center); ILC (innate lymphoid cell); ISC (intestinal stem cell); logFC (log fold change); LZ (light zone); PCA (principal component analysis); PEDV (porcine epidemic diarrhea virus); TA (transit amplifying)

To infer dynamics of PEDV infectivity in epithelial cells, pseudotime trajectories of epithelial cell transitionary pathways were constructed based on gene expression data (**Figure 4B-C**). Trajectory analysis indicated four transitional pathways derived from intestinal stem cell/transit amplifying (ISC/TA) cells, transitioning through early/intermediary cell states that ultimately polarized into mature enterocytes in cluster 3 (trajectory 1), mature enterocytes in cluster 38 (trajectory 2), goblet cells (trajectory 3), and *BEST4*+ enterocytes (trajectory 4) (**Figure 4D**). Assessment of trajectories 1 and 2 in PEDV samples demonstrated PEDV infection occurred at the terminus of each trajectory, with PEDV-infected cells arising around the same pseudotime value in both trajectories and reaching similar levels of PEDV RNA copies by the end of each trajectory (**Figure 4E; Supplementary Figure 5B**).

Trajectories 1 and 2 indicated mature enterocytes transitioned into distinct, transcriptionally-polarized states that were the primary targets of PEDV infection, and the two trajectories bifurcated from each other around the same pseudotime when PEDV infection increased (red dotted line in **Figure 4E**). We further investigated potential conversion between cells at the termini of the two trajectories by constructing another pseudotime trajectory containing only enterocyte cells occurring after the bifurcation of trajectories 1 and 2 (**Figure 4F; Supplementary Figure 5C**). The new trajectory inferred possible transition between cells of trajectories 1 and 2, and ends of the new trajectory still polarized towards cells found in only one of the two original trajectories (**Figure 4G**). We further compared the specific transcriptional profiles of polarized enterocyte states by identifying top DEGs (**Supplementary File 4**) between cells identified exclusively at the terminus of trajectory 1 versus trajectory 2 (green versus purple populations, respectively, in **Figure 4G**). Results indicated trajectory 1 polarized into mature enterocytes expressing genes associated with homeostatic functions in nutrient absorption and maintenance of barrier integrity [31–36] (**Figure 4H**), while trajectory 2 polarized into mature enterocytes expressing higher percentages of mitochondrial genes (**Figure 4I**), indicating cellular stress [37]. Thus, trajectory 1 gave rise to a population of homeostatic mature enterocytes, while trajectory 2 gave rise to a population of stressed mature enterocytes.

### Transcriptional reprogramming converges into a 190-gene signature shared amongst PEDV-infected and bystander enterocytes in homeostatic or stressed states

Both homeostatic and stressed mature enterocytes were primary targets of PEDV infection, yet how PEDV infection status affects enterocyte transcriptional profiles remained to be determined. DEGs were identified for homeostatic and stressed enterocytes (green and purple populations, respectively, in **Figure 4G**) between pairwise comparisons of infected cells (*PEDV*+ cells from PEDV animals), bystander cells (*PEDV*-cells from PEDV animals), and mock cells (cells from mock animals) (**Supplementary File 5**). In both homeostatic (**Figure 5A**) and stressed enterocytes (**Figure 5B**), more DEGs were identified between bystander or infected cells relative to mock cells than between bystander and infected cells.

**Figure 5.**
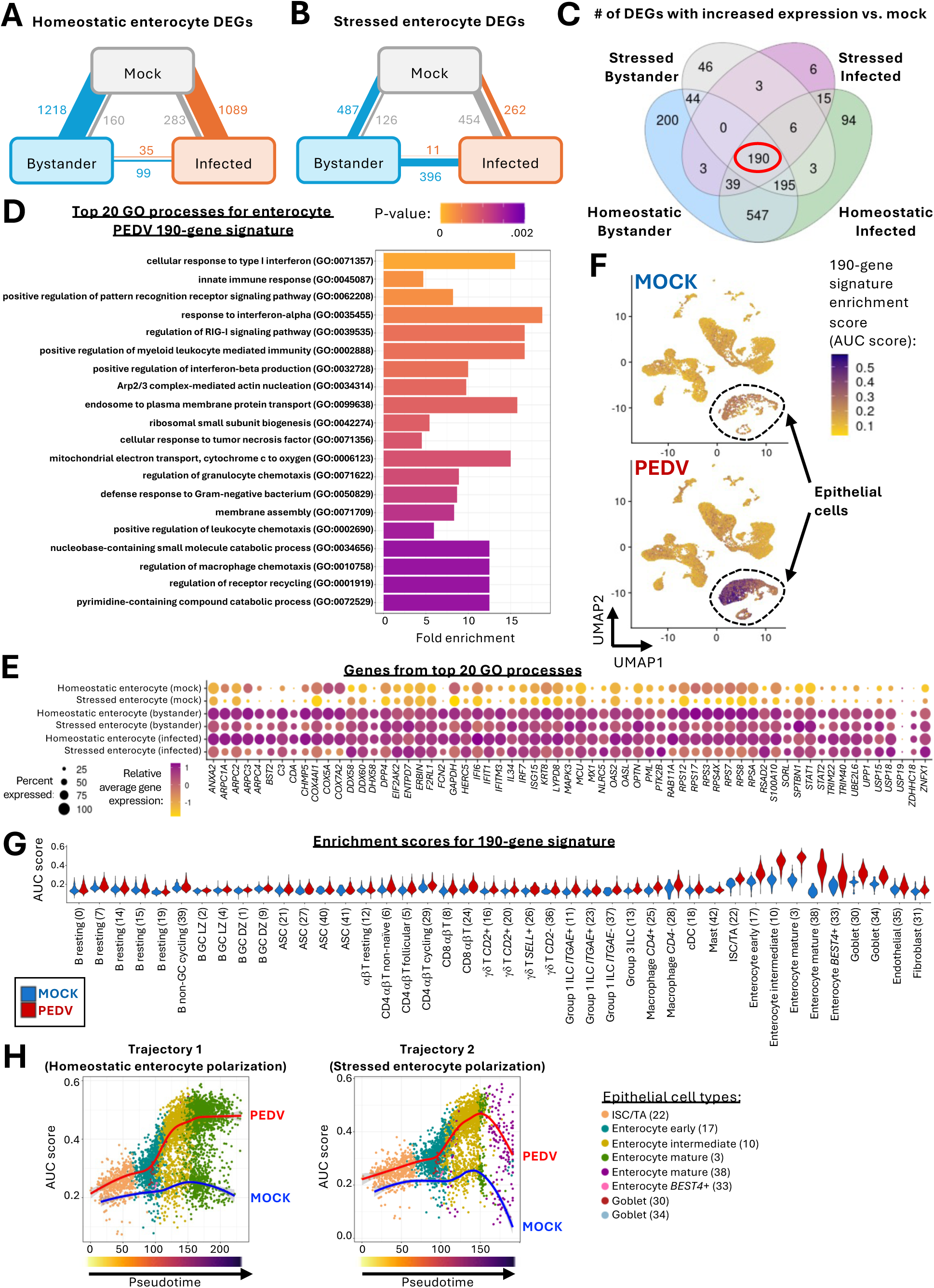
A conserved transcriptional program is initiated across enterocytes in response to PEDV infection. **(A-B)** Weighted graphs indicating the number of DEGs identified through pairwise comparisons between mock (grey), bystander (blue), and infected (orange) cells identified within total homeostatic enterocytes **(A)** or stressed enterocytes **(B)**. Lines indicate number of DEGs recovered by pairwise comparison between the two cell states each line connects. Line thickness correlates positively with the number of DEGs. Line color corresponds to the cell state increased expression was observed in. DEGs had corrected p-values <0.05, were expressed in at least 10% of one cell population being compared, and had a |logFC| >0.25. **(C)** Venn diagram showing the number of genes commonly identified as DEGs with increased expression in homeostatic bystander versus homeostatic mock enterocytes (blue), stressed bystander versus stressed mock enterocytes (grey), stressed infected versus stressed mock enterocytes (purple), and homeostatic infected versus homeostatic mock enterocytes (green). A total of 190 genes (circled in red) were identified as DEGs in all four comparisons. **(D)** Bar plot showing 20 GO processes (y-axis) with the lowest p-values (bar color) enriched from the 190-gene signature identified in **(C)**. Fold enrichment of GO processes is shown on the x-axis. A biological process was considered significant if it had a p-value <0.05 and had at least two detected genes contributing to the GO term. **(E)** Dot plot showing expression patterns in enterocyte subsets (y-axis) for a subset of genes (x-axis) included in the 190-gene signature identified in **(C)**. Displayed genes were associated with the 20 GO processes shown in **(D)**. Dot size corresponds to the percent of cells expressing a gene within each enterocyte subset. Dot fill color corresponds to the relative average expression value of a gene, for those cells expressing a gene within each enterocyte subset. **(F)** UMAP plot of cells shown in Figure 2A-D. Each dot represents one cell. The color of the dot corresponds to a gene set enrichment score (AUC score) for the 190-gene signature identified in **(C)**. **(G)** Violin plot of gene set enrichment scores (AUC scores, y-axis) for the 190-gene signature identified in **(C)** detected in cell clusters of mock (blue) or PEDV (red) samples (x-axis). Each dot represents one cell from a respective cluster. **(H)** Scatter plots showing gene set enrichment scores (AUC scores, y-axes) for the 190-gene signature identified in **(C)** across pseudotime values (x-axes) for trajectory 1 (left) and 2 (right) shown in (Figure 4C-E). Each dot represents one cell. Dot color corresponds to assignment of a cell to a cluster. A line of best fit shows gene set enrichment scores across pseudotime for PEDV samples (red) or mock samples (blue). Abbreviations: ASC (antibody-secreting cell); AUC (area under the curve); cDC (conventional dendritic cell); DEG (differentially expressed gene); DZ (dark zone); GC (germinal center); GO (Gene Ontology); ILC (innate lymphoid cell); ISC (intestinal stem cell); logFC (log fold change); LZ (light zone); PEDV (porcine epidemic diarrhea virus); TA (transit amplifying); UMAP (uniform manifold approximation and projection)

Of the DEGs with increased expression in bystander or infected relative to mock cells of respective homeostatic or stressed enterocytes, 190 genes were present in all comparisons (**Figure 5C; Supplementary File 6**). The conserved 190-gene signature included genes associated with antiviral and innate immune responses, such as pattern recognition receptor or type I interferon signaling responses (**Figure 5D-E; Supplementary File 7**). The 190-gene signature was further applied to additional cell clusters of PEDV versus mock samples, and the gene set was identified as enriched primarily in epithelial cells of PEDV samples (**Figure 5F)**. The highest gene signature enrichment scores were identified in intermediate and mature enterocyte clusters of PEDV samples (**Figure 5G**), and fitting of enrichment signature scores to mature enterocyte polarization trajectories (trajectories 1 and 2 from **Figure 4C-D**) demonstrated increasing differences in enrichment scores for the 190-gene signature in PEDV versus mock cells as enterocyte polarization (trajectory pseudotime) progressed (**Figure 5H**).

## DISCUSSION

The intestine is a complex tissue environment, with functionally unique compartmentalization, e.g. epithelium, lamina propria, Peyer’s patches, muscularis. In this work, we focused specifically on dynamics of transcriptional reprogramming for epithelial barrier defense and Peyer’s patch immune induction in response to PEDV. Recently weaned pigs were used to study PEDV infection dynamics, as the immediate post-weaning period introduces increased susceptibility to PEDV due to waning maternal immunity, an underdeveloped immune system, and increased weaning-associated stressors and antigen exposures [38]. We collected tissue samples of jejunum containing Peyer’s patches at 5dpi, when peak PEDV infection and initial induction of an adaptive immune response can be detected [39–41]. Results collectively demonstrate PEDV infection elicits widespread modulation of the intestinal cellular landscape, with diverse cell types adopting transcriptional programs promoting a cumulative antiviral response.

Peyer’s patches are unique immune inductive sites that directly sample luminal antigens due to positioning of organized lymphoid tissue in the intestinal submucosa, negating the need for lymphatic migration of APCs and promoting a more specialized mucosal immune response [23]. Although Peyer’s patches may be critical to eliciting localized protective immunity in the intestine, their importance as immune inductive sites against PEDV infection is underexplored. PEDV+ cells have been noted in Peyer’s patches as early as 2dpi [42], and Peyer’s patch follicular areas are enlarged by oral PEDV vaccination [43], indicating Peyer’s patches are exposed to and activate follicular immune processes in response to PEDV antigens. However, cellular interactions occurring in Peyer’s patches during PEDV infection need be further understood to promote favorable immune inductive processes protective against PEDV. In our work, signaling pathways associated with T cell-dependent B cell activation were identified as enriched in PEDV samples, indicating transcriptional promotion of processes where Tfh cells serve as central hubs for receiving signals derived from antigen-presenting macrophages, cDCs, and non-resting B cells. Tfh cells, non-resting B cells, and *CD4*+ macrophages have previously been shown to be primarily associated with Peyer’s patches within the pig small intestine [44, 45], indicating above described signaling interactions are likely associated with Peyer’s patch immune induction. Studies of immune induction in porcine Peyer’s patches have focused largely on cDCs as primary APCs [46] and have identified increased *in situ* cDC numbers and *in vitro* stimulatory capacity of cDCs due to PEDV antigen exposure [47]. Other recent work has identified functionally-unique macrophage subsets (*CD4+* or *CD4-*) that may be important APCs in the pig intestine [44], yet the role of these newly described cells in antiviral immunity has not been explored. In our work, both cDCs and macrophage subsets contributed to increased signaling associated with Tfh cell activation, and both may be important APCs for antiviral immunity that warrant further investigation. Overall, results indicate an important role of Peyer’s patches in orchestrating anti-PEDV immune responses, but the specificity and efficacy of such a response, and whether similar dynamics occur with other enteric viral infections, remains to be further evaluated. In future work, stronger emphasis may be placed on Peyer’s patches as a target for eliciting protective immunity to enteric viruses, including PEDV, and immune reactions occurring within Peyer’s patches may serve as readouts of localized immune induction that possibly proxy protection against viral infections. With recent developments in veterinary immunoreagent availability, *ex vivo* identification of Tfh cells [48], intestinal macrophage subsets [44], and functional B cell subsets [44, 49] may offer timely and important tools for further exploration of Peyer’s patch immune induction in pigs, including applications to both swine health and potential biomedical modeling [38, 44, 45, 50–53].

Previous reports indicate villus enterocytes are a primary target of PEDV infection [42, 54], yet the mechanisms by which diverse subsets of enterocytes (e.g. stressed versus homeostatic; infected versus bystander) craft antiviral transcriptional responses within a PEDV-infected tissue environment are undetermined. Our results indicate mature enterocytes reach two highly polarized cell states, homeostatic and stressed profiles, that are both primary targets of PEDV infection, and enterocytes may transition between homeostatic and stressed states. PEDV may play a role in initial enterocyte polarization to a stressed state or transition from homeostatic to stressed states, as PEDV induces epithelial cell stress [55]. Regardless, both homeostatic and stressed enterocytes were susceptible to PEDV infection, corresponding to likely locations of late-stage mature enterocytes in the apical villus epithelium [56], where most PEDV infections are documented *in situ* [42, 54] and were also detected in our work. Homeostatic and stressed enterocytes were further differentiated into infection states, including mock, bystander, and infected cells. PEDV caused widespread transcriptional reprogramming to both cells directly infected with virus and bystander cells exposed to an infected tissue microenvironment. While infected cells are subjected to direct effects of viral infection, uninfected bystander cells are likely modulated by environmental cues that are integral components of a coordinated response in the intestine. We identified a 190-gene signature that was conserved across actively infected and bystander enterocytes that were either homeostatic or stressed, indicating enterocytes responded to PEDV infection via conserved upregulation of a set of genes that was largely associated with antiviral responsiveness. The 190-gene signature was enriched in epithelial cells, with increasing enrichment detected in a differentiation stage-dependent manner that indicated highest expression of signature genes in more mature enterocytes. Interestingly, not all genes in the signature may be associated with protective responses against PEDV. Some genes within the 190-gene signature encode receptors directly targeted by PEDV for entry into host cells (*ANPEP* [57, 58]) or cofactors that facilitate PEDV invasion and replication (*DPP4* [59]). Other genes in the signature encode receptors identified as important cofactors for other coronaviruses, such as *PLAC8* for swine acute diarrhea syndrome coronavirus (SADS-CoV [60]) and *CDH17* for severe acute respiratory syndrome coronavirus 2 (SARS-CoV-2) [61]. Moreover, some signature genes are associated with regulation of intracellular processes critical for coronavirus assembly and trafficking (*ANXA2, ARPC3*, *ATP8B1* [62–64]), epithelial disruption (*MPP5* [65]), and suppression of host antiviral defenses (*CASP3, LYZ, TRIM40, USP15* [66–69]). Thus, the conserved enterocyte 190-gene signature may be indicative not only of antiviral transcriptional programs generated in response to PEDV infection, but some genes may also contribute to increased viral susceptibility and/or be exploited by PEDV. Genes identified in the conserved signature should be further explored to identify targets that can be strategically modulated to develop PEDV control strategies.

Recovered cell types and signaling interactions are in general consensus with results obtained from previous studies assessing the single-cell landscape in healthy pig intestinal tissues of similar age and rearing [44, 45, 70–73]. The single-cell transcriptional landscapes occurring during PEDV infection have been reported previously [74, 75], and our work has both commonalities and differences with previously reported results. All works identified a coordinated antiviral transcriptional response occurring across diverse subsets of immune and epithelial cells in response to PEDV infection [74, 75]. Similar to our findings, one work also identified enterocytes as the primary target of PEDV infection, though other cell types were also infected [75]. However, distinctions in PEDV challenge isolate (virulent Chinese versus United States PEDV strains), inoculation dosage (2 or 5 mL ∼10^6^ TCID_50_/mL versus 2 mL 10^5^ TCID_50_/mL), pig age (three days old versus four weeks old), infection time course (24 or 60 hours post-inoculation versus 5dpi), sample type (jejunum without versus with Peyer’s patches), sequencing sample replicates (single pooled treatment samples versus individual treatment replicates), and dataset size (12,774 or 19,612 versus 40,522 total cells) provided distinct biological scenarios and datasets that were being analyzed in the previous reports [74, 75] versus our work, respectively.

Altogether, this work encompasses a high-resolution transcriptional investigation of host-pathogen interactions occurring during PEDV infection. Results emphasize the coordination of an antiviral immune response, including global reprogramming of diverse cell types across the complex intestinal landscape. Newly defined dynamics pertaining to immune induction and epithelial defense strategies supply important information that can be applied to better understand mechanisms of PEDV pathogenesis and identify novel preventative or therapeutic targets to control PEDV.

## MATERIALS AND METHODS

Extended materials and methods are available in **Supplementary File 8**. Briefly, PEDV non-S-INDEL isolate USA/NC49469/2013 [76, 77] was grown to a final titer of 10^5^ TCID_50_/mL. Approximately 4-week-old pigs were inoculated orally with 2 mL PEDV (n=5) or mock (n=3) inoculum. Daily rectal swabbing and diarrhea scores were collected until euthanasia at 5dpi. At necropsy, the most proximally-located Peyer’s patch was grossly identified and collected for histopathology, viral quantification, and cell isolations. Histologic evaluation included blinded pathology examination of hematoxylin and eosin (H&E)-stained tissue sections using metrics for atrophic enteritis scoring available in **Supplementary File 9.** *In situ* assessment of PEDV localization was assessed via immunohistochemistry staining for PEDV nucleoprotein. Viral quantification was performed via RT-qPCR to detect viral copies of RNA isolated from rectal swabs and collected tissue. Cells were isolated from tissue to use as input for scRNA-seq.

scRNA-seq was performed using the Chromium Single-cell 3’ Kit v3 (10X Genomics). Data quality control and processing were performed similar to previous workflows [44, 45, 70, 78, 79]. Using Cell Ranger v8.0.1 (10X Genomics), reads were aligned to a *Sus scrofa* 11.1 genome incorporating the PEDV USA/NC/2013/49469 complete reference genome (GenBank KM975737.1) as an additional gene sequence. Ambient RNA was removed using SoupX v1.6.2 [80], and doublets were estimated using scDblFinder v1.16.0 [81]. Low quality cells (doublets, ≥15% mitochondrial reads, ≤250 genes, ≤3000 unique molecular identifiers [UMIs]) were removed. Data were further processed using standard SCTransform (SCT) normalization, canonical-correlation analysis (CCA)-based data integration, uniform manifold approximation and projection (UMAP) dimensionality reduction, and clustering procedures with Seurat v5.1.0 [82]. Cell clusters were annotated based on canonical gene expression patterns described previously [44, 45, 70]. Pseudobulk analysis of overall sample compositions was performed using edgeR v4.0.16 [83] as previously described [45]. DGE analysis was performed with a fast Wilcoxon rank sum test and area under the receiver operating characteristic curve (auROC) analysis based on Gaussian approximations using presto v1.0.0 [84]. Enriched Gene Ontology (GO) biological processes were identified using the bottom-up elim method and Fisher’s exact test with topGO v2.54.0 [85] and biomaRt v2.58.2 [86] as previously described [44, 45]. Cell signaling networks were inferred using CellChat v2.1.2 [87] as previously described [44]. Pseudotime trajectory analysis was performed using Slingshot v2.10.0 [88] similar to previous work [44]. Gene set enrichment analysis was performed using AUCell v1.24.0 [89] as previously described [78, 79].

## DATA AVAILABILITY

Raw sequencing data is deposited at SRA and will be made publicly available under BioProject PRJNA1263541 upon publication. Data will be made available on the Broad Single-cell Portal for interactive online query upon publication. Materials for interactive query using Loupe Browser (10X Genomics; .cloupe file formats), processed data objects available for computational investigation (.h5Seurat and .rds file formats), and selected source data will be made available at Ag Data Commons upon publication. Scripts used for data analysis and visualization are available at https://github.com/SwiVi/scRNAseq_PEDV_JejunalPeyersPatches.

## ACKNOWLEDGEMENTS

This work was funded by USDA-ARS CRIS project #5030-32000-230-000-D and an appointment to the Agricultural Research Service (ARS) Research Participation Program administered by the Oak Ridge Institute for Science and Education (ORISE) through an interagency agreement between the U.S. Department of Energy (DOE) and the United States Department of Agriculture (USDA). This research used resources provided by the SCINet project of the USDA ARS project numbers 0201-88888-003-000D and 0201-88888-002-000D. ORISE is managed by Oak Ridge Associated Universities (ORAU) under DOE contract number DE-SC0014664. All opinions expressed in this paper are the authors’ and do not necessarily reflect the policies and views of USDA, ARS, DOE, or ORAU/ORISE. Mention of trade names or products is for information purposes only and does not imply endorsement by the USDA. USDA is an equal opportunity employer and provider.

We are grateful to the following for their contributions: Sarah Anderson and Colin Stoy for laboratory assistance; Lauren Tidgren Hanson, Andrew Von Weber, and London Harris for sample collection assistance; Adrienne Shircliff for histology assistance; the National Animal Disease Center Animal Resources Unit for animal care; the Iowa State University DNA Facility for scRNA-seq library preparation and sequencing; Dr. Darrell Bayles for initial data management; Dr. Daniel Nielsen for computational assistance; Drs. Luis Gimenez-Lirola, Meghan Wymore Brand, and Bryan Kaplan for subject matter input; Dr. Tavis Anderson for manuscript review.

Authors contributed to this work as described in the following. **Jayne E. Wiarda**: conceptualization, methodology, software, investigation, formal analysis, resources, data curation, visualization, supervision, project administration, writing – original draft, writing – review & editing. **Bailey Arruda**: investigation, formal analysis, visualization, writing – review & editing. **Eraldo L. Zanella**: methodology, investigation, writing – review & editing. **Hanjun Kim**: formal analysis, writing – review & editing. **Samantha J. Hau**: investigation, writing – review & editing. **Jianqiang Zhang**: resources, writing – review & editing. **Alexandra C. Buckley**: methodology, investigation, resources, writing – review & editing.

## SUPPLEMENTARY FIGURES

**Supplementary Figure 1.**
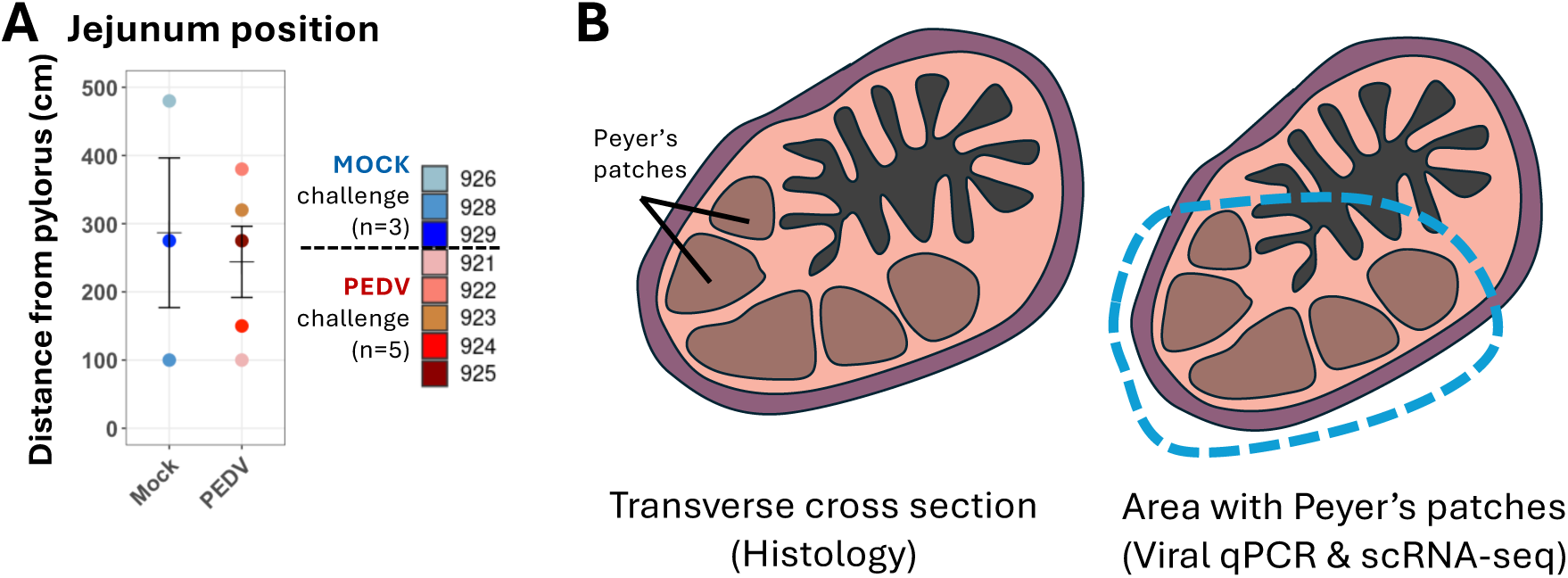
Collection of jejunum containing Peyer’s patches mock and PEDV-inoculated pigs. **(A)** Distance from the pylorus (y-axis) where each sample of proximal-most jejunal Peyer’s patch was identified and collected from mock (blue) or PEDV (red) pigs (x-axis) at 5dpi. Bar intervals indicate mean and standard error of the mean. **(B)** Representative images showing transverse cross sections of jejunum containing Peyer’s patches that were taken for histology versus tissue sections that were further dissected to contain only regions with Peyer’s patches that were taken for viral quantification and scRNA-seq. Abbreviations: cm (centimeters); dpi (days post-inoculation); PEDV (porcine epidemic diarrhea virus); qPCR (quantitative polymerase chain reaction); scRNA-seq (single-cell RNA sequencing)

**Supplementary Figure 2.**
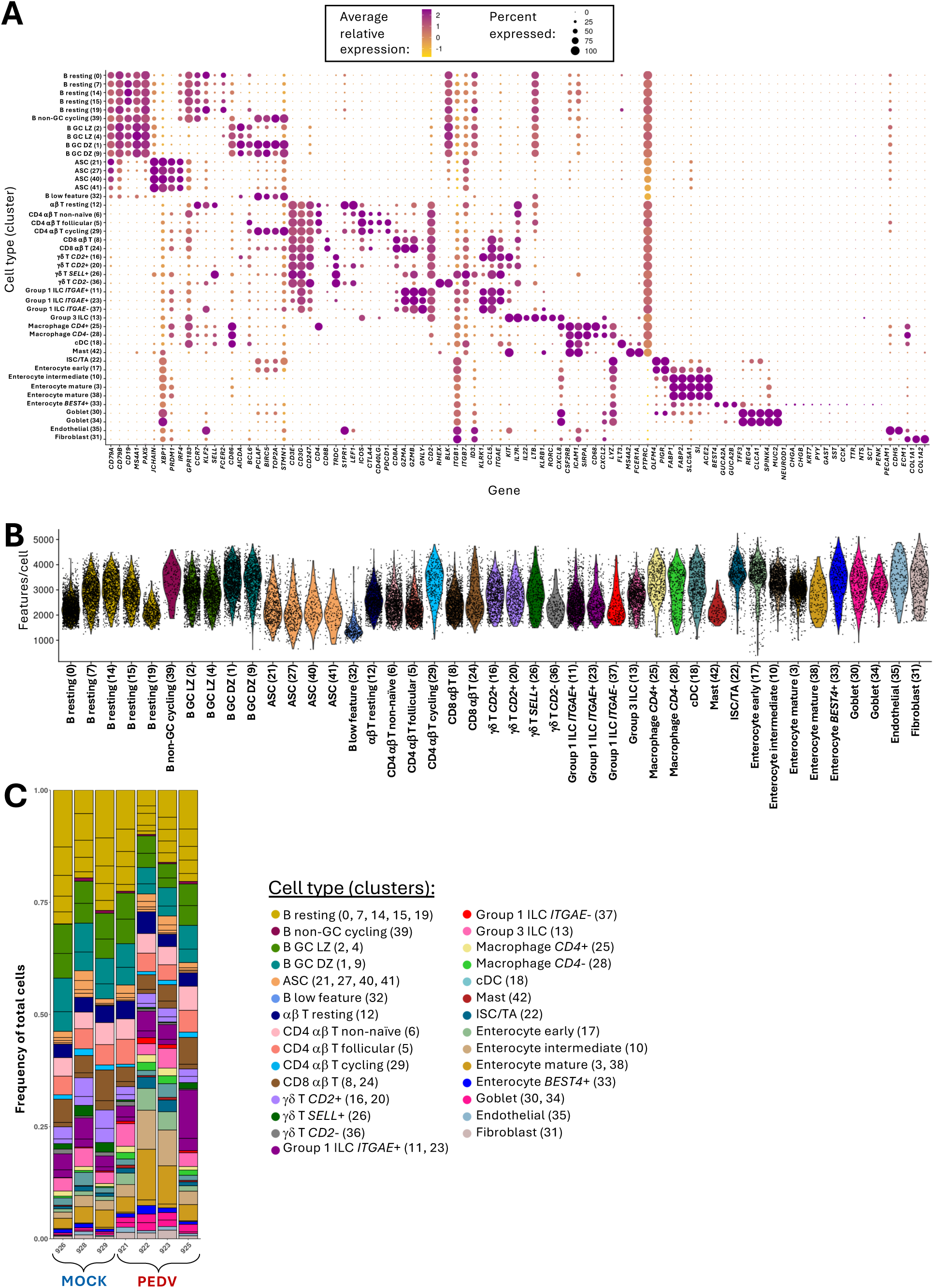
Cell types recovered from jejunum containing Peyer’s patches via scRNA-seq. **(A)** Dot plot showing expression patterns in cell clusters (y-axis) for canonical genes (x-axis) used to annotated clusters into cell types. Dot size corresponds to the percent of cells expressing a gene within each cluster. Dot fill color corresponds to the relative average expression value of a gene, for those cells expressing a gene within each cluster. **(B)** Violin plot of the number of features (y-axis) detected in cell clusters (x-axis). Each dot represents one cell from a respective cluster. **(C)** Stacked bar plots showing the frequency of cells (y-axis) belonging to each cell cluster (fill colors) across animal samples (x-axis). Abbreviations: ASC (antibody-secreting cell); cDC (conventional dendritic cell); DZ (dark zone); GC (germinal center); ILC (innate lymphoid cell); ISC (intestinal stem cell); LZ (light zone); PEDV (porcine epidemic diarrhea virus); scRNA-seq (single-cell RNA sequencing); TA (transit amplifying)

**Supplementary Figure 3.**
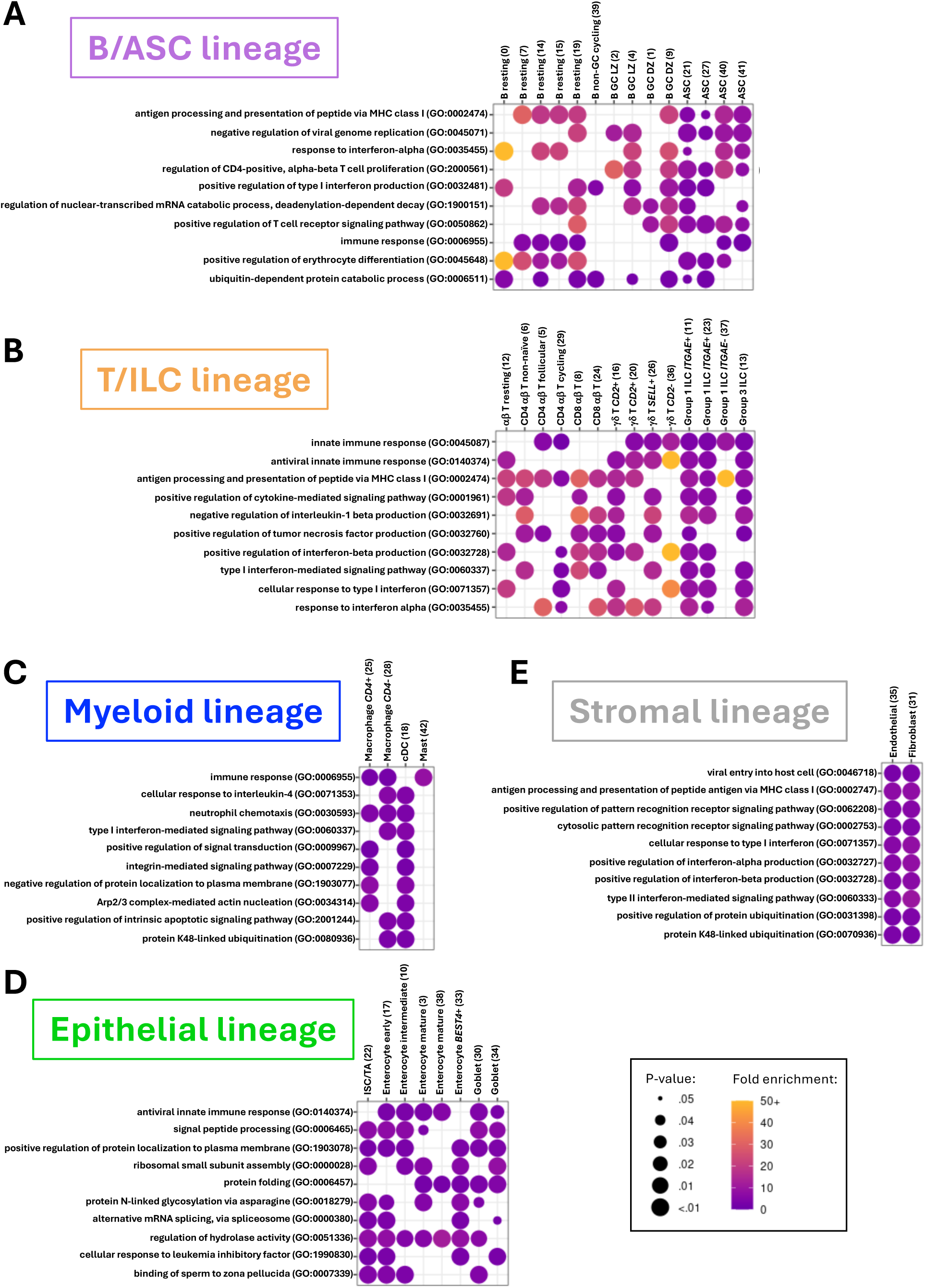
Biological processes enriched across diverse jejunal cell types and lineages during PEDV infection. **(A-E)** Dot plot of the top 10 GO terms (y-axes) significantly enriched in PEDV relative to mock cell clusters (x-axes) within B/ASC **(A)**, T/ILC **(B)**, myeloid **(C)**, epithelial **(D)**, and stromal **(E)** lineages. Dot color indicates fold enrichment; dot size indicates p-value. The top 10 biological processes were selected as those found to be significantly enriched in the majority of cell clusters within a lineage that also had the smallest median p-values. A biological process was considered significant if it had a p-value <0.05 and had at least two detected genes contributing to the GO term. Abbreviations: ASC (antibody-secreting cell); cDC (conventional dendritic cell); DZ (dark zone); GC (germinal center); GO (Gene Ontology); ILC (innate lymphoid cell); ISC (intestinal stem cell); LZ (light zone); PEDV (porcine epidemic diarrhea virus); TA (transit amplifying)

**Supplementary Figure 4.**
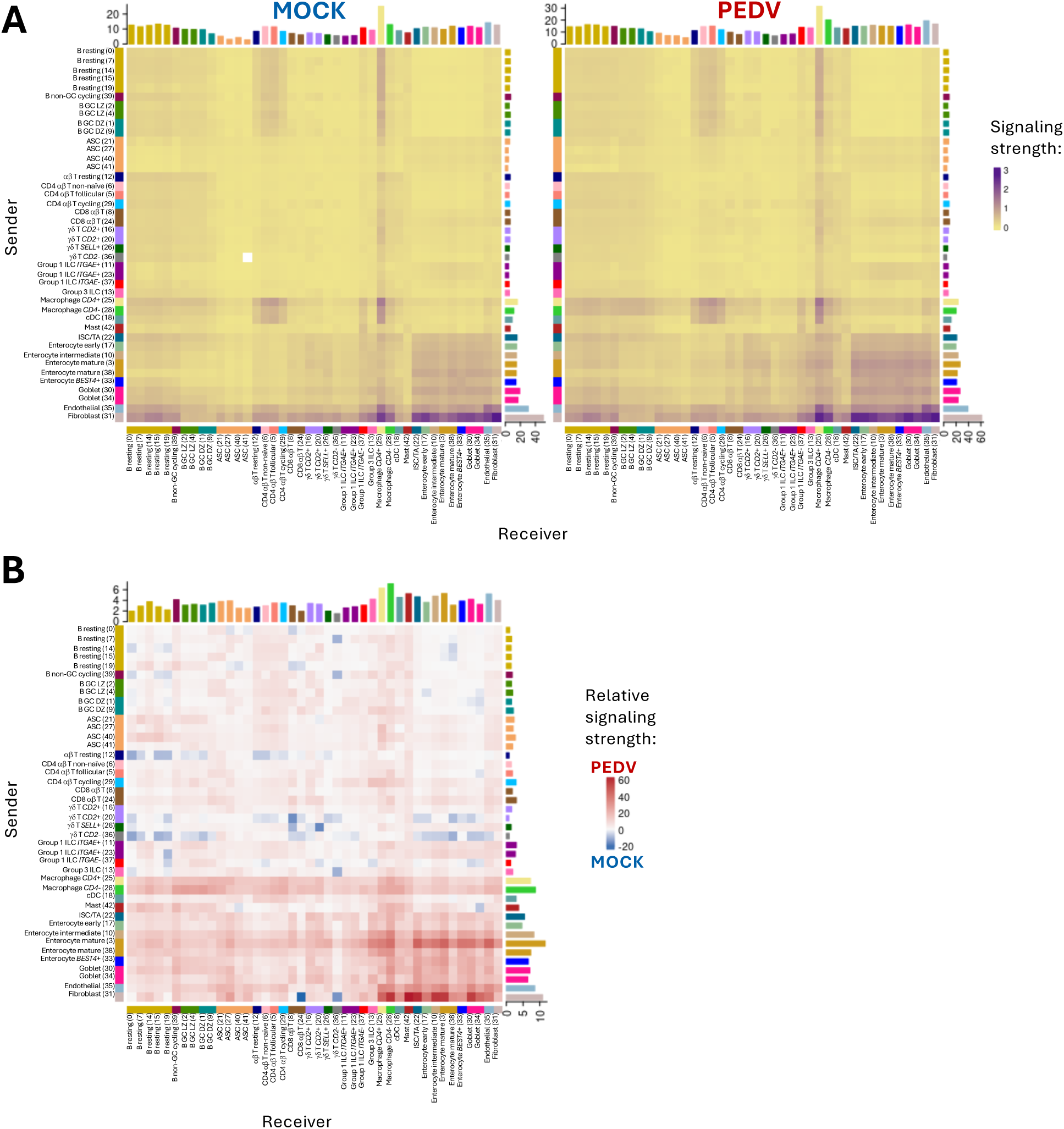
Cumulative signaling strength networks across cell types of mock or PEDV cells. **(A)** Heatmaps from mock (left) or PEDV (right) scRNA-seq datasets where cumulative cell signaling strength (fill color) is indicated for pairwise interaction comparisons of all cell clusters as senders (y-axes) or receivers (x-axes) of signaling interactions. **(B)** Heatmap showing the relative differences in signaling strengths for mock (blue) or PEDV (red) samples based on values shown in **(A)**. Abbreviations: ASC (antibody-secreting cell); cDC (conventional dendritic cell); DZ (dark zone); GC (germinal center); ILC (innate lymphoid cell); ISC (intestinal stem cell); LZ (light zone); PEDV (porcine epidemic diarrhea virus); scRNA-seq (single-cell RNA sequencing); TA (transit amplifying)

**Supplementary Figure 5.**
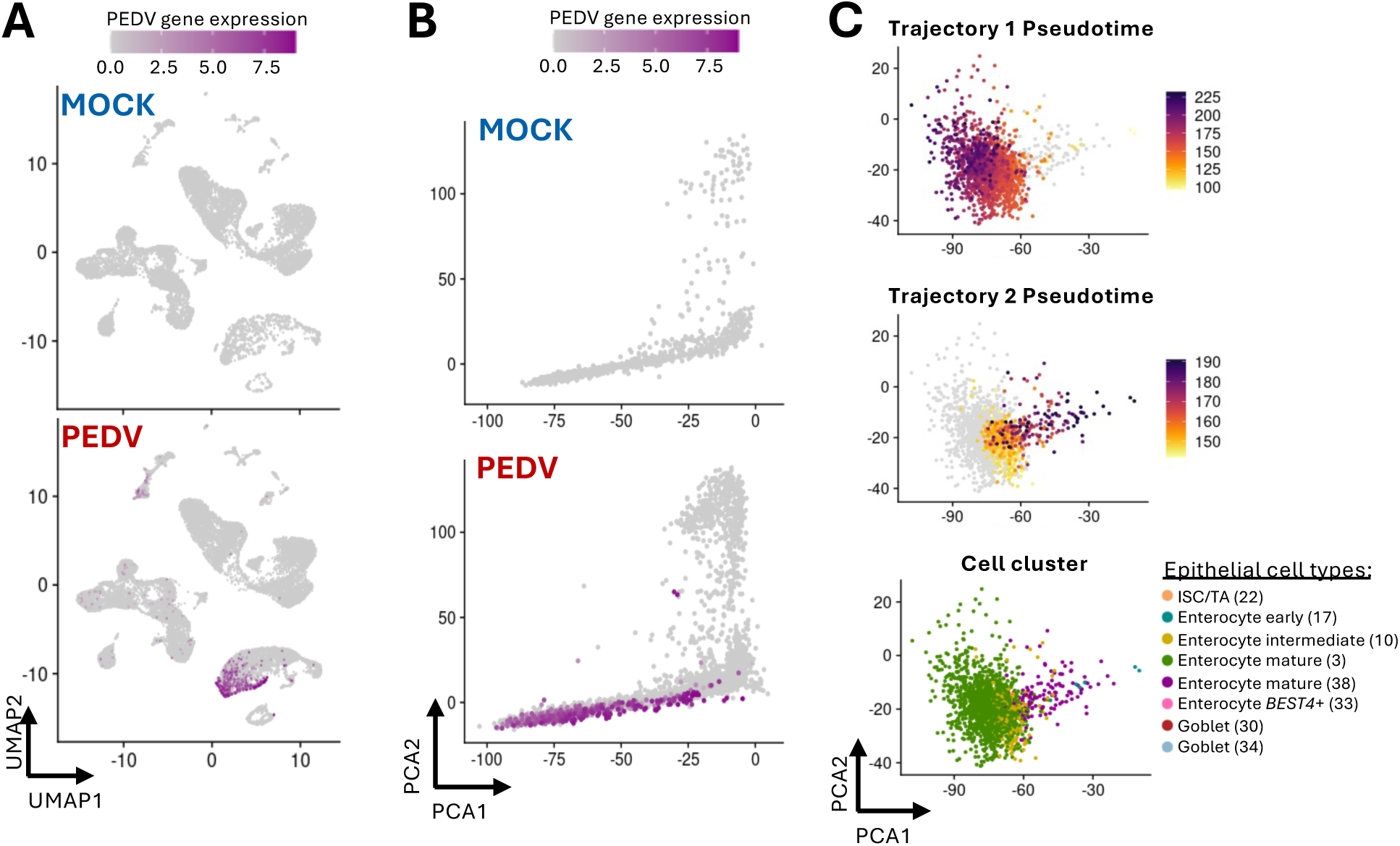
PEDV detection and trajectory analysis of enterocytes. **(A-B)** UMAP **(A)** or PCA **(B)** plot of cells shown in Figure 2A-D **(A)** or Figure 4B-C **(B)**. Each dot represents one cell. The color of the dot corresponds to expression level for PEDV RNA. Cells from mock samples are shown in the top panel; cells from PEDV samples are shown in the bottom panel. **(C)** PCA plot of cells shown Figure 4F. Each dot represents one cell. The color of the dot corresponds to pseudotime values of cells from trajectory 1 from Figure 4C (top), trajectory 2 from Figure 4C (middle), or cell cluster (bottom). Abbreviations: ISC (intestinal stem cell); PCA (principal component analysis); PEDV (porcine epidemic diarrhea virus); TA (transit amplifying); UMAP (uniform manifold approximation and projection)

