## SupplementaryFile8 for "Porcine epidemic diarrhea virus infection promotes Peyer’s patch immune induction and epithelial defense via single-cell transcriptional reprogramming"

### Supplementary File 8: Materials and methods (extended)

Extended materials and methods from:

"Porcine epidemic diarrhea virus promotes Peyer's patch immune induction and epithelial defense via single-cell transcriptional reprogramming" by Jayne E. Wiarda, Bailey Arruda, Eraldo L. Zanella, Hanjun Kim, Samantha J. Hau, Jianqiang Zhang, and Alexandra C. Buckley

#### Animals and experimental overview

Eight conventional, mixed-breed pigs were obtained from a commercial farrowing site that was negative for porcine epidemic diarrhea virus (PEDV). Pigs were weaned at approximately 3 weeks of age, transported to the National Animal Disease Center (NADC) in Ames, Iowa, placed into biosafety level (BSL)-2 housing, and acclimated for approximately 1 week before experimental inoculation. Five pigs in one room received 2 mL PEDV inoculum orally, while three pigs in a second room received 2 mL of mock cell culture medium orally. PEDV inoculum was prepared by passaging PEDV non-S-INDEL isolate USA/NC49469/2013 [1, 2] eight times in Vero cells, with a final grown titer of $10^5$ $TCID_{50}$/mL used for inoculation. Diarrhea scores and rectal swabs were obtained from animals pre-challenge (0 days post-inoculation [dpi]) and daily thereafter until necropsy at 5dpi. At necropsy, animals were humanely euthanized via intravenous administration of barbiturate (Fatal Plus, Vortech Pharmaceuticals) followed by collection of tissue samples. All animal work was performed according to an Animal Care and Use Protocol (ARS-24-1197) approved by the NADC Institutional Animal Care and Use Committee.

#### Diarrhea scoring

Pigs were observed daily for signs of diarrhea and received a score based on the following categorizations: (0) normal feces, (1) loose feces, (2) mild diarrhea, (3) voluminous, watery diarrhea.

#### Rectal swab collection

Rectal swabs were collected with a sterile polyester-tipped applicator (Puritan Medical Products) immersed in 3 mL serum-free minimum essential medium (MEM) and stored at −80°C until testing.

#### Tissue collection

The most proximally-located Peyer's patch was collected by working distally from the pylorus until a Peyer's patch was grossly identified. The distance of each Peyer's patch from the pylorus was recorded for each animal. A longitudinal section containing the Peyer's patch was excised, and the Peyer's patch was further cut in half to create two comparable longitudinal pieces. One half was collected into 10% neutral-buffered formalin (NBF) for later histology. The second half was further dissected to exclude regions not containing Peyer's patches (similar to non-Peyer's patch samples in previous work [3]). A small section of the second half of tissue was placed into RNAlater (Thermo Fisher AM7021) for later RNA isolation/viral quantification, and the remainder was placed into RPMI 1640 (Gibco 11875-085) containing 5% fetal calf serum (FCS) (Seradigm, Avantor 1600-50) for cell isolation.

#### Viral quantification

Quantification of PEDV nucleic acid via reverse transcription quantitative polymerase chain reaction (RT-qPCR) was performed as previously described [4]. For rectal swabs, RNA extraction was performed using a MagMAX Viral RNA Isolation Kit (Applied Biosystems AMB18365) following the manufacturer's recommendations for fecal samples. For tissue stored in RNAlater, tissue was homogenized in TRI Reagent (Thermo Fisher AM9738) using M-tubes (Miltenyi Biotec 130-093-236) on a GentleMACS OctoDissociater (Miltenyi Biotec 130-134-029), protocol RNA 01_01. The MagMAX-96 for Microarrays Total RNA Isolation Kit (Applied Biosystems AM1839) was used for RNA isolation following the spin procedure. RNA was quantified using the

Bioanalyzer RNA 6000 Nano Kit (Agilent 5067-1511) on a 2100 Bioanalyzer system (Agilent). Following RNA extractions, 5 μl (rectal swab RNA) or 500 ng (tissue) of isolated RNA were added to Path-ID multiplex one-step RT-PCR master mix (Applied Biosystems 4442137) to make a 25 μl reaction, and RT-qPCR was performed on an ABI 7500 Fast instrument (ThermoFisher). PEDV genome copy numbers were calculated based on a plasmid overlapping the target region. Duplicate well values for each sample were averaged together.

#### Immunohistochemistry

Longitudinal halves of jejunum containing Peyer's patches that had been collected into 10% NBF at necropsy were processed into formalin-fixed, paraffin-embedded (FFPE) tissue blocks and sectioned onto slides as previously described [5].

Slides were baked, deparaffinized, and rehydrated, followed by sequential application of the following: Proteinase K (Dako S3020) 3 min room temperature (RT), BLOXALL (Vector SP-6000) 10 min RT, 2.5% horse serum (component of Vector MP-6402) 30 min RT, anti-PEDV nucleoprotein antibody clone 3D6 (Immunology Consultants Laboratory MPEDV-5A-3D6; stock concentration 5.4 mg/mL; diluted 1:10,000 in 1% bovine serum albumin [BSA; Sigma A0281] in phosphate-buffered saline [PBS; made in-house, pH 7.2]) overnight 4ºC, horse anti-mouse IgG peroxidase polymer (component of Vector MP-6402) 30 min RT, 3,3'-diaminobenzadine (DAB; Vector SK-4103) 7 min RT, Gill's Hematoxylin I (StatLab HXGHE) 1 min RT. Slides were washed twice with PBS containing 0.05% Tween 20 (Sigma P1379) 2 min RT between all steps except serum and anti-PEDV nucleoprotein antibody incubations. Slides were next rinsed, dehydrated, and coverslipped as previously described [5].

#### Pathology scoring & assessment

Tissue sections stained with hematoxylin and eosin (H&E) of jejunum containing Peyer's patches were assessed by a pathologist that was blinded to PEDV versus mock sample treatments. Pathology scoring was performed on a 0 to 4 scale to define signs of atrophic enteritis, including villus blunting, villus fusion, crypt elongation, and epithelial attenuation as summarized in **Supplementary File 9**.

After conducting blinded scoring, tissue sections of jejunum containing Peyer's patches that were stained for PEDV nucleoprotein via immunohistochemistry were assessed by the same pathologist to identify tissues with positive versus negative PEDV staining. Within positively stained tissues, PEDV nucleoprotein distributions were analyzed and qualitatively reported.

#### Cell isolation

Collected jejunal tissue containing Peyer's patches was transported to the lab in 5% FCS RPMI at RT. Approximately 1.25 g tissue was weighed and used for further processing. All starting solutions were equilibrated to RT unless otherwise specified. For all shaking steps, tubes containing tissue and reagent were placed at a 45° angle and shaken at 150 rpm on a Max Q 4000 Shaking Incubator (Thermo Scientific SHKE4000). Weighed tissue was placed into 50 mL conicals containing 10 mL RPMI with 1 mM EDTA (Invitrogen AM9260G) and 1 mM dithiothretol (Invitrogen 15508013) and incubated 37°C 10 min shaking. Tissues were transferred to C-tubes (Miltenyi Biotec 130-096-334) containing 7 mL RPMI with 0.2 U/mL Liberase TM (Roche 0540427001) and 30 ug/mL DNase (Qiagen 79254). Tissues in C-tubes were mechanically dissociated with the B_01 protocol on a GentleMACS OctoDissociater (Miltenyi Biotec 130-134-029), incubated 37°C 30 min shaking, and dissociated again with the B_01 protocol. 7 mL chilled 10% FCS RPMI was added to each C-tube, and homogenate was poured over a 500-micron filter (PluriSelect 43-50500-01) into an empty 50 mL conical and placed on ice. The collection was centrifuged 300xg 4°C 10 min. Supernatants were poured off, and cell pellets were resuspended in 15 mL ACK lysis buffer (made in-house) for 3 min RT. 30 mL 5% FCS RPMI was added to each tube, and again tubes were centrifuged and supernatants poured off. Cell pellets were resuspended in volumes (10-30 mL depending on pellet size) of 5% FCS RPMI and placed on ice. Collections were poured over a 40-micron filter (PluriSelect 43-57040-01) into an empty 50 mL conical and placed on ice. Cell concentrations and viability were assessed using the Muse Count & Viability Kit (Cytek Biosciences MCH600103) read on a Muse Cell Analyzer (Luminex) as previously described [3, 6, 7]. Using the obtained concentration readings, a total of $1x10^6$ total cells from each sample were diluted in 5% FCS RPMI to a total volume of 1 mL and transported to the Iowa State University DNA Facility (~15 min) for immediate single-cell RNA sequencing (scRNA-seq).

#### Single-cell RNA sequencing

##### ***Partitioning, library preparation, and sequencing***

Cells were submitted to the Iowa State University DNA facility for partitioning and library preparation using the Chromium Single-cell 3' Kit v3 (10X Genomics) as previously described [3, 6-9]. 10,000 cells were targeted for partitioning from each sample. cDNA libraries were sequenced across two lanes of a NovaSeq 6000 S1 flow cell (Illumina) as previously described [3, 6-9]. One sample library (pig #928) had a lower recovered sequencing depth per cell, and deeper sequencing was performed using ~20% capacity across two lanes of an additional NovaSeq 6000 SP flow cell. Reads from each sequencing run were merged together in subsequent steps after verifying adequate quality of reads yielded from each run.

#### Single-cell RNA sequencing data analyses

Data analysis was performed using interactive RStudio sessions (R v4.3.2) [10] within the Ceres high performance computing infrastructure available from SciNet, available through the Agricultural Research Service (ARS). Scripts used for each step of data analyses are available as detailed in the main text's data availability statement. The data availability statement also indicates availability of additional data objects, including finalized data objects used for computational analyses and interactive data objects that can be accessed for non-computational data queries.

##### *Data quality control and processing*

Data quality control and processing were performed similar to previous workflows [3, 6-9]. Default parameters were used for all processes unless otherwise specified. fastQC v0.12.1 [11] was used to assess sequence quality and deem sufficient for downstream applications. Cell Ranger v8.0.1 (10X Genomics) was used to align reads to the *Sus scrofa* 11.1 genome with a custom annotation file used previously [7]. To also annotate and count polyA viral sequences for the PEDV challenge strain, the pig genome and annotation file were further modified to append the PEDV USA/NC/2013/49469 complete reference genome (GenBank KM975737.1) as a single exon sequence in the pig reference genome and annotation file as per manufacturer's instructions (10X Genomics). Cell Ranger output files were used to perform ambient RNA calculation and removal with SoupX v1.6.2 [12] followed by doublet estimation with scDblFinder v1.16.0 [13]. Cells that had ≥15% mitochondrial reads, ≤250 genes detected, ≤3000 unique molecular identifiers (UMIs) detected, and/or were predicted as doublets were removed. Genes with sum zero raw counts in all remaining cells were also removed. Seurat v5.1.0 [14] was further used to process data using standard methods for data normalization with SCTransform (SCT), data integration with canonical correlation analysis (CCA) methods, and calculation of dimensional reductions (principal component analysis [PCA], CCA-based PCA, t-distributed stochastic neighbor embedding [t-SNE], uniform manifold approximation and projection [UMAP]).

##### *Cell type annotation*

Seurat v5.1.0 [14] was used to cluster cells. Shared nearest neighbors were calculated with the integrated CCA data reduction, and a clustering resolution of 1.5 was used to create clusters. Canonical gene expression patterns described previously [3, 6-8] were used to annotate clusters. Several clusters with similar expression of key canonical genes were merged into single annotations.

B cells and antibody-secreting cells

B cell clusters (clusters 0, 1, 2, 4, 7, 9, 14, 15, 19, 39) expressed genes encoding components of the B cell receptor (*CD79A, CD79B, CD19, MS4A1*) and the B lineage transcription factor, *PAX5,* while antibody-secreting cell (ASC) clusters (clusters 21, 27, 40, 41) were identified by expression of ASC markers *JCHAIN, XBP1, PRDM1, IRF4* [3, 7, 8]. Resting B cell clusters (clusters 0, 7, 14, 15, 19) had high expression of markers associated with naïve or memory B cells, including *GPR183, CCR7, KLF2, SELL, FCER2*, while non-resting B cell clusters had higher expression of activation marker *CD86* [3, 7]*.* Of the non-resting B cell clusters (clusters 1, 2, 4, 9, 39), cluster 39 was identified as a non-germinal center (GC) cycling B cell cluster due to lack of expression for genes associated with GCs (*AICDA, BCL6*) and high expression of cell cycle genes (*PCLAF, BIRC5, TOP2A, STMN*) [3, 7]. Remaining non-resting B cell clusters (clusters 1, 2, 4, 9) expressed genes associated with GC localization and activation (*BCL6, AICDA*) and were differentiated into clusters of GC light zone (LZ) B cells (clusters 2, 4) and GC dark zone (DZ) B cells (clusters 1, 9) by weak or strong expression of cell cycle genes (*PCLAF, BIRC5, TOP2A, STMN*), respectively [7].

T cells

T cell clusters (clusters 5, 6, 8, 12, 16, 20, 24, 26, 29, 36) had strong expression of genes associated with the T cell receptor complex, including *CD3E, CD3G, CD247*, and were further called as CD4 αβ T cells, CD8 αβ T cells, or γδ T cells by expression of *CD4, CD8B,* or *TRDC*, respectively [3, 7, 8]. Cluster 12 included cells predominately expressing *CD4* but also some *CD8B*-expressing cells, indicating a cluster of CD4 and CD8 αβ T cells. Cluster 12 further had high expression of resting T cell markers, including *CCR7, KLF2, SELL, S1PR1, LEF1*, and was thus annotated as resting αβ T cells [3, 7, 8]. Three additional clusters with *CD4* expression (clusters 5, 6, 29) were classified as non-resting subsets of CD4 αβ T cells based on expression of activation markers, *ICOS, CTLA4, CD40LG* [3, 7, 8]. Cluster 29 had high expression of cell cycle genes (*PCLAF, BRIC5, TOP2A, STMN*) and was called as CD4 αβ T cycling cells [3, 7]. Higher versus lower expression of follicular T helper cell marker *PDCD1* was used to call clusters 5 and 6 as follicular and non-naïve CD4 αβ T cell clusters, respectively [3, 7]. Two clusters of CD8 αβ T cells (clusters 8, 24) expressed *CD8B* but also *CD8A* and cytotoxicity-associated genes (*GZMA, GZMB, GNLY*) that confirmed identities [3, 7]. γδ T cell clusters (clusters 16, 20, 26, 36) were first divided into CD2 lineages, where lack of *CD2* expression in cluster 36 was used to identify *CD2*- γδ T cells [3, 7, 8]. *CD2*- γδ T cells also had strong expression of *RHEX, BLK*, which are proposed to also be genes with strong expression in *CD2*- T cells [3, 7, 8]. Of the remaining γδ T cells with strong *CD2* expression, one cluster also had high expression of *SELL, ITGB1, ITGB7, ID3*, which are genes used to describe a recently-discovered subset of *SELL*+ γδ T cells in pig intestine [3, 7]. Remaining γδ T cell clusters 16 and 20 were classified as *CD2*+ γδ T cells.

<u>Innate lymphoid cells</u>

Innate lymphoid cells (ILCs) share close transcriptional similarities to many T cell subsets and have high expression of *CD2* but low/no expression of *CD3E* [3, 7], and such criteria was used to identify clusters of ILCs in our data (clusters 11, 13, 23, 37). Three ILC clusters (clusters 11, 23, 37) expressed *CD8A*, *KLRK1, CD3G, CCL5, GZMA, GZMB, GNLY*, which are markers recently established to be associated with group 1 ILCs in pig intestine, including potential natural killer (NK) cells and ILC1 cells [3, 7]. *ITGAE* (encoding CD103) has been proposed as an important marker to differentiate resident versus non-resident ILCs, as well as intraepithelial lymphocytes [3], and group 1 ILC clusters were thus divided into *ITGAE-* group 1 ILCs (cluster 37) and *ITGAE*+ group 1 ILCs (clusters 11, 23). The last ILC cluster (cluster 13) lacked expression of CD3 T cell receptor accessory molecule genes (*CD3E, CD3G, CD247*) and had high expression of genes including *KIT, IL7R, IL22, LTB, KLRB1, RORC, CXCL8* that are highly expressed in group 3 ILCs, including both lymphoid tissue inducer (LTi) cells and ILC3 cells [3, 7]. Cluster 13 was thus annotated as group 3 ILCs.

Macrophages, dendritic cells, and mast cells

Myeloid lineage cells (clusters 18, 25, 28, 42) expressed markers including *CSF2RB* and *ICAM1* [3, 7, 8]. Macrophage clusters (clusters 25, 28) were identified by strong expression of *SIRPA, CD68, CXCL2, LYZ* [3, 7, 8]. *CD4* expression was used to classify macrophages as *CD4*+ macrophages (cluster 25) or *CD4*- macrophages (cluster 28) based on recent discovery of these macrophage subsets in pig intestine [7]. Cluster 18 was annotated as conventional dendritic cells (cDCs) based on high expression of *FLT3* [3, 7, 8]. Similar to previous reports [8], cDCs also expressed *SIRPA* but at levels lower than macrophages. Mast cells were identified in cluster 42 by expression of *MS4A2, FCER1A* [3, 7].

Epithelial and stromal cells

Remaining clusters (clusters 3, 10, 17, 22, 30, 31, 33, 34, 35, 38) included non-immune cells that lacked expression of the pan-leukocyte marker, *PTPRC* [3, 6, 7]. Clusters 17 and 22 had strong expression of epithelial crypt cell markers, *OLFM4, PIGR*, *LYZ* and cell cycle genes (*PCLAF, BRIC5, TOP2A, STMN*), indicating a location in intestinal epithelial crypts [6, 7]. Cluster 17 also expressed markers for nutrient transport and absorption (*FABP1, FAPB2, SLC5A1, SI, ACE2*), while cluster 22 did not. Thus, cluster 22 was annotated as intestinal stem cells (ISCs)/transit amplifying (TA) cells, while cluster 17 was annotated as early enterocytes [6, 7]. Clusters 10, 3, and 38 also had strong expression of markers for nutrient absorption/transport, indicating enterocyte identities [6, 7]. However, cluster 10 still had low-level expression of crypt markers, leading to differentiation of intermediate enterocytes in cluster 10 and mature enterocytes in clusters 3 and 38 [6, 7]. Cluster 33 was annotated as *BEST4*+ enterocytes due to expressing genes including *BEST4*, *GUCA2A, GUCA2B,* which are recognized to be expressed in newly-recognized subsets of *BEST4* enterocytes present in pig intestine [6, 7]. Clusters 30 and 34 were called as goblet cells due to expression of canonical goblet cell markers including *TFF3, REG4, CLCA1, SPINK4, MUC2* [6, 7]. Unlike previous works [6, 7], an enteroendocrine (EE) cell cluster was not identified in the dataset. Canonical EE cell genes including *NEUROD1, CHGA, CHGB, KRT7, PYY, GAST, SST, CCK, TTR, NTS, SCT, PENK* [6, 7] were queried, with no cluster having strong expression. It was noted that a very small percentage of cells in the *BEST4*+ enterocyte cluster (cluster 33) had low-level expression of some EE cell genes, suggesting a miniscule number of cells in the cluster could potentially be EE cells, but the numbers were ultimately too low to create an analyzable EE cell cluster in this dataset. Cluster 35 was identified as endothelial cells by expression of *PECAM1* and *CDH5*, and cluster 31 was identified as fibroblasts by expression of *ECM1, COL1A1, COL1A2* [7].

##### *Pseudobulk analysis*

Pseudobulk analysis of overall sample compositions was performed using SCT-normalized reads stored in the Seurat object after cell and gene filtering. Analysis was performed using edgeR v4.0.16 [15] to obtain a multidimensional scaling plot of data as previously described [3].

##### *Differential gene expression analyses*

Differential gene expression (DGE) analysis was performed using a fast Wilcoxon rank sum test and area under the receiver operating characteristic curve (auROC) analysis based on Gaussian approximations of SCT-normalized counts data with the presto v1.0.0 package [16]. Further filtering was applied to identify differentially expressed genes (DEGs) as having an adjusted p-value <0.05, absolute value of log fold-change (|logFC|) >0.25, and expression in at least 10% of one cell group being compared.

##### *Biological process enrichment analysis*

Enriched Gene Ontology (GO) biological processes were identified as previously described [3, 7] using the bottom-up elim method and Fisher’s exact test with topGO v2.54.0 [17] and biomaRt v2.58.2 [18]. Lists of DEGs with increased expression for a cell group were used as input gene lists for analysis. GO terms with p-values <0.05 and at least 2 genes contributing to the term were considered significantly enriched biological processes.

##### *Cell signaling network analysis*

CellChat v2.1.2 [19] was applied to identify cell signaling networks inferred from gene expression data. Gene counts were humanized as previously described [7] in order to apply the human cell interaction database available through CellChat. All cell signaling types (cell-cell contact, secreted, extracellular matrix [ECM]-receptor, non-protein signaling) were applied to define signaling networks. Communication probabilities were calculated using the truncatedMean method with a trim value of 0.1. A merged CellChat object was created by merging separate CellChat objects of PEDV and mock samples. Default parameters were used to identify interaction strengths and enriched signaling pathways based on relative information flows.

##### *Pseudotime trajectory analysis*

A data subset containing only epithelial lineage cells was created, with new dimensionality reductions and integrations being calculated for the data subset, similar to previous work [7, 9]. Pseudotime trajectories were constructed using Slingshot v2.10.0 [20]. Trajectories were first created using all epithelial cells, where ISC/TA cluster 22 was specified as a starting cluster, and grouping resolution for trajectory construction was specified at the cell cluster level. Using tradeSeq v1.16.0 [20], a generalized additive model (GAM) was fit to the recovered trajectory data using seven knots. The optimal number of knots to use was determined based on selection criteria previously used [7]. The fourth knot (pseudotime=148.796) was used as a cutoff pseudotime value to create another subset of data based on expression of PEDV RNA increasing after this knot value in trajectory 1 and 2. The new data subset included only cells occurring after a pseudotime value of 148.796 in trajectory 1 or 2. The new data subset was also re-processed as outlined above. A trajectory for this subset was drawn with no specification of grouping factor or starting point due to only two clusters present. Due to lack of specification for a starting point, the trajectory direction was undefined and shown in a subjectively selected direction that indicated potential bidirectionality.

##### *Classification of enterocyte states: homeostatic versus stressed or bystander versus infected*

The data subset of cells occurring after pseudotime values of 148.796 from trajectory 1 or 2 (see pseudotime trajectory analysis methods above) was further analyzed to identify cells only occurring in trajectory 1 or 2. DGE analysis was performed to compare gene expression between cells of the data subset that only occurred in trajectory 1 versus 2. Based on DGE results, cells occurring after trajectory 1 pseudotime value of 148.796 and not found in trajectory 2 were annotated as homeostatic enterocytes, and cells occurring after trajectory 2 pseudotime value of 148.796 and not found in trajectory 1 were annotated as stressed enterocytes. Homeostatic or stressed enterocytes within PEDV samples were further assessed for expression of PEDV RNA within SCT-normalized counts data. If a cell contained any PEDV RNA, it was classified as an infected cell. If a cell had zero counts for PEDV RNA but still belonged to a PEDV-infected animal sample, it was classified as a bystander cell. All homeostatic or stressed enterocytes derived from mock animals were classified as mock cells.

##### *Gene set enrichment analysis*

Pairwise DGE comparisons (see DGE methods above) were made between mock, infected, and bystander cells within each of homeostatic or stressed enterocytes. DEGs with increased expression in homeostatic infected versus mock, homeostatic bystander versus mock, stressed infected versus mock, and stressed bystander versus mock enterocytes were identified, and a total of 190 genes were commonly identified in all four comparisons. The 190 genes were utilized as a gene signature for gene set enrichment analysis (GSEA).

GSEA was performed using AUCell v1.24.0 [21] as previously described [8, 9]. Gene counts were ranked by raw expression values, and area under the curve (AUC) enrichment values were calculated from the top 5% of ranked genes overlapping with the 190-gene signature.
